# Hypoxia Exacerbates Kir2.1 Channel Dysfunction in an Andersen-Tawil Syndrome Variant Through a SUMO-Dependent Mechanism

**DOI:** 10.1101/2025.04.25.650734

**Authors:** Aishwarya Chandrashekar, Yu Xu, Xinyi Ma, Anne K. Yauch, Elizabeth Scholl, Yuchen Yang, Kirin D. Gada, Takeharu Kawano, Meng Cui, Leigh D. Plant

## Abstract

**Background:** Andersen-Tawil Syndrome type 1 (ATS1) is a multisystem channelopathy that predisposes patients to ventricular dysrhythmias and increases the risk of sudden cardiac death. ATS1 arises from loss-of-function mutations in Kir2.1, the inward rectifying potassium channel responsible for most of *I_K1_* in ventricular cardiomyocytes. *I_K1_* is suppressed by SUMOylation, a post-translational modification upregulated in hypoxia, a known proarrhythmic stimulus. We investigated whether current from the ATS1-linked variant Kir2.1-R67Q is inhibited by hypoxia and whether this suppression can be reversed by pharmacological inhibition of the SUMO pathway.

**Methods:** We used patch-clamp recording to measure *I_K1_* and Kir2.1 currents under acute hypoxia, with and without the SUMO pathway inhibitor TAK-981. To quantify SUMOylation stoichiometry, we applied single molecule photobleaching. A multidisciplinary approach combining electrophysiology, molecular modeling, and optogenetic phosphoinositide was used to measure the impact of Kir2.1- R67Q and SUMOylation on channel interactions with phosphatidylinositol 4,5-bisphosphate (PIP_2_), a required gating cofactor.

**Results:** Kir2.1 can be modified by up to two SUMO proteins attached to diagonally opposite subunits, with each SUMOylation event reducing current by ∼20%. Heterozygouse channels containing two R67Q subunits were more susceptible to hypoxic suppression than wild type. TAK-981 blocked hypoxic inhibition of *I_K1_* in ventricular cardiomyocytes and abolished Kir2.1 SUMOylation. In cells expressing Kir2.1-R67Q, TAK-981 significantly increased currents and mitigated hypoxic suppression. Computational modeling and optogenetic dephosphorylation revealed that both the R67Q mutation and converge to disrupt Kir2.1- PIP_2_ interactions, producing synergistic inhibition of channel function.

**Conclusions:** Hypoxia-induced SUMOylation and the R67Q mutation synergistically suppress Kir2.1 activity by impairing channel-PIP_2_ interactions. TAK-981 restores *I_K1_* by preventing SUMOylation under hypoxic conditions and enhancing current through Kir2.1-R67Q channels. These findings support a two-hit model of arrhythmogenesis in ATS1 and identify SUMO pathway inhibition as a potential therapeutic strategy to reduce arrhythmic risk in affected patients.

## Introduction

Andersen-Tawil Syndrome type 1 (ATS1) is a rare, autosomal dominant, multisystem disorder caused by loss-of-function mutations in *KCNJ2*, the gene that encodes for the inward-rectifying K⁺ channel, Kir2.1. Even though ATS1 is typically a heterozygous condition, patients may present with periodic paralysis, distinctive dysmorphic features, and severe cardiac dysrhythmia, including prolonged QT intervals (LQT), augmented U waves, and complex premature ventricular contractions (PVCs), often accompanied by tachyarrhythmia ^1–7^. Although ATS1 predisposes patients to arrhythmia, the overall incidence of adverse cardiac events is surprisingly low ^8–11^. However, as with other channelopathies, the risk of adverse events, including sudden cardiac arrest, is significantly increased by environmental triggers such as sympathetic stimulation or drug side effects. This pattern aligns with the two-hit hypothesis of arrhythmogenesis ^8–13^.

Kir2.1 channels conduct the majority of the *I_K1_* current in the myocardium. ATS1 variants reduce *I_K1_*, leading to cardiomyocyte depolarization and prolonged action potential duration due to slowed phase 3 repolarization ^14–16^. Kir2.1 function relies on interactions with the membrane phospholipid phosphatidylinositol 4,5-bisphosphate (PIP₂), a critical cofactor required for channel gating. As a result, small molecule regulators, post-translational modifications (PTMs), and disease mutations that disrupt Kir2.1-PIP_2_ interactions can reduce channel activity. Several ATS1-associated variants, such as Kir2.1-R67Q are proposed to exert their effects through this mechanism ^1, 15, 17–19^. Conversely, strategies aimed at enhancing *I_K1_* have been proposed as preventative approaches in patients predisposed to specific arrhythmic phenotypes ^14^.

We recently reported that SUMOylation of Kir2.1 reduces *I_K1_* in cardiomyocytes by decreasing the efficacy of PIP₂-mediated channel activation ^15^. SUMOylation is a multi-step enzymatic PTM in which one of three SUMO isoforms is covalently coupled to the ε-amine group of specific lysine residues on target proteins. The process begins with SUMO activation by the E1-enzyme, SAE1, followed by transfer to the E2 conjugating enzyme, Ubc9, which recognizes the SUMOylation motif, lysine 49 in the case of Kir2.1 ^20–22^. While basal SUMOylation of Kir2.1 is low, it increases rapidly under hypoxic conditions, leading to proarrhythmic suppression of *I_K1_* ^11, 15, 20^. Given the critical role of hypoxia-induced regulation of Kir2.1 in cardiac function, we tested whether blocking SUMOylation could increase the activity of ATS1-variant Kir2.1 channels.

Here, we demonstrate that hypoxia and the ATS1-lined Kir2.1-R67Q mutation act synergistically to suppress Kir2.1 activity by disrupting PIP_2_-dependent channel gating. Using a combination of single-channel biophysics, computational modeling, and cardiomyocyte electrophysiology, we show that SUMOylation underlies this suppression and that inhibition of the SUMO pathway with TAK-981 restores *I_K1_* by stabilizing Kir2.1–PIP_2_ interactions. These findings define a mechanistic basis for hypoxia-induced arrhythmia susceptibility in ATS1 and support SUMOylation of Kir2.1 as a modifiable therapeutic target in genetically vulnerable patients ^12, 13^.

## Materials & Methods

### Reagents

Purified SUMO1 and SENP1 were purchased from R & D Systems. The monoclonal antibody against Kir2.1 (RRID: AB_11000720) was purchased from Neuromab and the polyclonal antibody for SUMO1 [Y299] (ab32058) was from Abcam. TAK-981 was from MedChemExpress, 2-D08 was from Selleck Chem, and N106 was from Tocris. The components of the Proximity Ligation Assay (DUO92101) and all other chemical reagents were purchased from Sigma-Aldrich.

### Molecular biology and Biochemistry

SUMO1 (GenBank: NM_003352.8), Ubc9 (GenBank: NM_003345.5), SENP1 (GenBank: NM_001267594.2) and Kir2.1 (NM_017296.1) were handled in pMAX, a CMV-based in-house vector^15^ for patch-clamp of HEK293T cells or to generate cRNA for subsequent two-electrode voltage-clamp recording from *Xenopus* oocytes. The mTFP1-tagged Kir2 constructs were in pMAX. Sequences encoding eYFP or mTFP1 were inserted at the N terminus of SUMO1 or the Kir2.1 variants, respectively. Mutations were introduced with Quikchange (Agilent) and verified by sequencing (Psomagen, Inc). CIBN-CAAX and mCherry-CRY2-5-ptase_OCRL_ were kind gifts from the De Camilli lab and are characterized in (Idevall-Hagren et al., 2012).

### Cell culture

Rat ventricular cardiomyocytes from neonatal Wistar rats were purchased from Lonza Biosciences. Cells were seeded at 300,000/ cm^2^ on 1% fibronectin coated coverslips or culture dishes (Thermo Fisher Scientific) and were maintained according to the manufactures instructions in Bullet complete medium (Lonza) supplemented with 200 µM bromo-uridine to inhibit the proliferation of cardiac fibroblasts. Human embryonic kidney (HEK293T) cells were acquired from American Type Culture Collection (ATCC) and were maintained in Dulbecco’s modified Eagle’s medium (ATCC) supplemented with 100 units/ml penicillin, 100 μg/ml streptomycin, and 10% (vol/vol) fetal bovine serum. All cells were incubated in a 37 °C, humified incubator supplemented with 5% CO_2_. Where indicated, cells were cultured at 7% O_2_ in a hypoxia-capable tri-gas incubator (Thermo). All cardiomyocytes studied by patch-clamp were identified by their beating phenotype. HEK293T cells were transfected using polyethylenimine solution (1 mg/ml) at a ratio of 6 µl per 1 µg of plasmid DNA, 24-30 hours prior to experiments. For patch-clamp, FRET, and control experiments, we transfected 1 µg of each plasmid DNA, as indicated. Cells for study were identified by expression of a fluorescence marker, typically eGFP for patch-clamp studies, or mCherry for optogenetic studies.

### Donor-decay FRET

Donor-decay time-course Förster Resonance Energy Transfer (FRET) was performed using an Olympus inverted epi-fluorescence microscope. HEK293T cells were transfected with the subunits indicated 24-30 hours prior to an experiment and studied in a solution comprising, in mM: NaCl 130, KCl 4, MgCl_2_ 1.2, CaCl_2_ 2, HEPES 10, pH was adjusted to 7.4 with NaOH. Regions of interest for study were first identified by YFP fluorescence using a 495/ 20 nm excitation filter and a 540/ 30 nm emission filter (Chroma). To assess donor-decay, mTFP1 was subjected to continuous excitation from a broad-spectrum LED (Excelitas) through a 436/ 20 nm filter using a 20x objective lens (Olympus). The output at the sample was measured at 50 mW/ cm^2^ by a light meter (ThorLabs). The emission was collected through a 480/ 40 nm bandpass filter (Chroma). Images were captured every 3 seconds using a sCMOS camera (Hamamatsu) controlled by µ-manager software ^23^. The exposure time was 300 ms. Images were saved as stack files and processed in ImageJ ^24^. Briefly, the mean fluorescence intensity of discrete areas of membrane, 3-5 nm in length was plotted against time and fit with a mono-exponential decay function to obtain the time constant for decay (Tau). 3-5 regions of interest from 24-35 cells were studied, generating 70-150 Taus per condition.

### Proximity ligation Duolink Assay

RVCM were fixed by 4% paraformaldehyde (PFA) and permeabilized by 0.2% Triton-X100. Primary antibodies for Kir2.1 (1:200) and SUMO1 (Y299, 1:250) were diluted in phosphate buffered saline and applied to the fixed cells at 4 °C and incubated overnight. A PLA assay kit (Sigma Aldrich, DUO92101) was used according to the manufacturer’s instructions. For negative control groups, antibodies against Kir channels were not used. Confocal pictures were captured on a Zeiss LSM880 microscope and analyzed by ZenBlue software (Zeiss) and ImageJ. Briefly, 600 nm confocal slices were used to capture the top of the cells in a field of view, including the plasma membrane and the upper portion of the nucleus. Fluorescence signals indicating ligation between SUMO1 and Kir2.1 were collected through a Texas Red filter set (Zeiss, set 31) and nuclei were captured using DAPI. Images were subjected to thresholding to remove signals below the background image intensity. The number of cells in each region of interest was determined by counting DAPI stained nuclei. Experiments were repeated at least three times to obtain overall PLA interactions as a function of the number of nuclei. Each field of view contains 20-30 nuclei. The image data shown are from a field of representative cells that is pseudo colored so that the nucleus is red and the PLA interactions are green for clarity. We previously confirmed the specificity of the Kir2.1 antibodies in our own hands by Western blot ^15^.

### Two-electrode voltage-clamp

Plasmid DNAs of Kir2.1, SUMO1, Ubc9, and SENP1 were linearized prior to *in vitro* transcription. Capped RNAs were transcribed using mMessage mMachine T7 transcription kit (Thermo Fisher Scientific). *Xenopus* oocytes were surgically extracted in accord with an IACUC protocol at Northeastern University. Oocytes were then dissociated and defolliculated by collagenase treatment and microinjected with 50 nl of a RNAse/ DNAse free water solution per oocyte containing the required cRNAs: we injected 1 ng channel, 5 ng SUMO1, 1 ng Ubc9, and 1 ng SENP1, as indicated, oocytes were incubated for 1-2 days at 17 °C before currents were recorded. Electrodes were pulled using a Flaming-Brown micropipette puller (Sutter Instruments) and were filled with 3 M KCl in 1.5% (w/v) agarose to give resistances between 0.5 and 1.0 MΩ. The oocytes were bathed in ND96 recording solution comprising, in mM: KCl 2, NaCl 96, MgCl_2_ 1, and HEPES 5, buffered to pH 7.4 with KOH. Where indicated, Kir2.1 currents were assessed in a high-K^+^ recording solution comprising, in mM: KCl 96, NaCl 2, MgCl_2_ 1, and HEPES 5, buffered to pH 7.4 with KOH. Currents were measured from whole oocytes using an OC-725D amplifier (Warner Instruments), controlled via a USB interface (National Instruments) and were recorded using WinWCP software (University of Strathclyde). To study Kir2 channels, oocytes were held at 0 mV, and currents were assessed by 160-ms ramps from -80 to +80 mV that were repeated every second. At the end of the study, currents were blocked by 10 mM BaCl_2_. The Ba^2+^-sensitive component of the current was analyzed. Between 4 and 12 oocytes from different *Xenopus* frogs were studied per experiment.

### Whole-cell patch clamp recording

Whole-cell currents were recorded with a Tecella Pico-2 amplifier (Tecella) controlled using WinWCP software (Spider, University of Strathclyde). Currents were acquired through a low-pass Bessel filter at 2 kHz and were digitized at 10 kHz. Patch pipettes were fabricated from borosilicate glass (Clark Kent), using a vertical puller (Narishige) and had a resistance of 2.5–4 MΩ when filled with the intracellular buffers described below. To study *I_K1_* in cardiomyocytes, the recording pipettes were filled with a solution comprising, in mM: K-aspartate 100, KCl 40, MgCl_2_ 5, EGTA 5, Na_2_ATP 5, and HEPES 5, adjusted to pH 7.2 (KOH). Cardiomyocytes were identified morphologically and by their beating behavior. Cardiac fibroblasts had a distinct morphology, did not beat, and did not observed to pass *I_K1_* current when they were studied by whole-cell patch-clamp recording (data not shown). The cells were initially bathed in a solution comprising, in mM: NaCl 140, KCl 5, MgCl_2_ 1, mM CaCl_2_ 1.8, glucose 10, and HEPES 5, adjusted to pH 7.4 (NaOH). To patch-clamp beating cells, the recording pipette was brought close to the cell of interest and prior to making contact, 10 µM nitrendipine was added to the recording solution to block Ca^2+^ flux through Ca_V_ channels. Following breakthrough into whole-cell mode, cardiomyocytes were allowed to stabilize for 2 mins, then *I_K1_* current was assessed by holding at 0 mV then stepping to +40 mV for 50 ms before a ramp to -120 mV at a rate of 1 mV/ ms. The magnitude of *I_K1_* was then assessed over 50 ms at -120 mV before the cell returned to the holding potential. The physiological recording buffer was replaced with a high-K^+^ buffer comprising, in mM: NaCl 81, KCl 64, MgCl_2_ 1, CaCl_2_ 1.8, glucose 10, and HEPES 5, adjusted to pH 7.4 (NaOH). Bath perfusion was via a multichannel gravity-driven perfusion manifold (Warner). This recording protocol, combined with the use of nitrendipine and the high K^+^ solution isolated *I_K1_* in the native cells ^19^. HEK293T cells heterologously expressing Kir2.1 channels were studied using an intracellular buffer comprising, in mM: KCl 140, MgCl_2_ 2, EGTA 1, Na_2_ATP 5 and HEPES 5, adjusted to pH 7.2 with KOH. Following breakthrough into whole-cell mode, the cells were allowed to stabilize for 2 mins then currents were assessed by ramps from -80 to +80 mV that were repeated at 1 Hz. The external physiological recording buffer comprised, in mM: NaCl 135, KCl 5, MgCl_2_ 1.2, CaCl_2_ 1.5, glucose 8, and HEPES 10, adjusted to pH 7.4 with NaOH, and transitioned to a high-K+ buffer comprising, in mM: NaCl 5, KCl 135, MgCl_2_ 1.2, CaCl_2_ 1.5, glucose 8, and HEPES 10, adjusted to pH 7.4 with KOH. For both native *I_K1_* and heterologous Kir2 channel studies, the Ba^2+^-sensitive component of the current was analyzed and determined by perfusing 5mM BaCl_2_ in the high-K+ buffer at the end of each experiment. Cells for study were selected based on GFP or mCherry expression using an epifluorescence microscope (Olympus). Rat cardiomyocytes and HEK293T cells had a mean whole-cell capacitance of 34 ± 5 pF and 10 ± 8 pF respectively; series resistance was typically <10 MΩ, and the voltage-error of 3 mV was not compensated.

### Generation and delivery of hypoxic perfusate

Acute hypoxia was achieved by switching the perfusate with one that had been bubbled with nitrogen for at least 30 minutes prior to the experiment. Oxygen tension was measured at the cell by a calibrated oxygen probe (Ocean Insight); solution exchange occurred in less than 20s via gas impermeable Tygon tubing. For patch-clamp studies, cells were initially studied in a quasi-physiological recording buffer, or a high-K^+^ buffer equilibrated to ambient O_2_ (21%) or 7% O_2_, as indicated. The cells were then exposed to a buffer equilibrated to 2% O_2_ (acute hypoxia) for the time indicated. For the PLA study, cells were exposed to hypoxia for 5-mins before being immediately fixed in 4% paraformaldehyde. All hypoxic solutions were equilibrated, measured, and perfused at room temperature (∼20 °C).

### Molecular Dynamics (MD) simulations

The protonation states of the titratable residues in the Kir2.1 channel and its mutants were calculated at pH = 7.4 via the use of the H++ server (http://biophysics.cs.vt.edu/) ^25^. The Kir2.1-PIP_2_, Kir2.1_R67Q-PIP_2_ or Kir2.1_R67Q_K49Q-PIP_2_ complexes were inserted into a simulated lipid bilayer composed of POPC:POPE:POPS:cholesterol (25:5:5:1) ^26^ and a water box using the CHARMM-GUI Membrane Builder webserver (http://www.charmm-gui.org) ^27^. Potassium chloride (150 mM) as well as neutralizing counter ions were applied to the systems. The total atom numbers are 248,273; 248,257; and 248,245 for the Kir2.1-PIP_2_, Kir2.1_R67Q-PIP_2_, and Kir2.1_R67Q_K49Q-PIP_2_, respectively. The PMEMD.CUDA program of AMBER 18 was used to conduct MD simulations ^28^. The Amber ff14SB, lipid17 and TIP3P force fields were used for the channels, lipids, and water, respectively. The parameters of PIP_2_ were generated using the general AMBER force field by the Antechamber module of AmberTools 17 and using the partial charge determined via restrained electrostatic potential charge-fitting scheme by ab initio quantum chemistry at the HF/6-31G* level ^29, 30^. PIP_2_ binding conformations were obtained from the Kir2.2/PIP_2_ crystal structure (PDB: 3SPI) by superimposition with Kir2.1. Coordinate files and system topology were established using the tleap module of Amber. The systems were energetically minimized by 500 steps (with position restraint of 500 kcal/mol/Å^2^) followed by 2,000 steps (without position restraint) using the steepest descent algorithm. Heat was then applied to the systems to drive the temperature from 0 to 303 K using Langevin dynamics with a collision frequency of 1 ps^-1^. The channel complexes were position-restrained using an initial constant force of 500 kcal/mol/Å^2^ during the heating process, subsequently diminished to 10 kcal/mol/Å^2^, allowing the lipid and water molecules free movement. Before the MD simulations, the systems underwent 5 ns of equilibration. Then, a total of 100 ns of MD simulations were conducted using hydrogen mass repartitioning and a time step of 4 fs. The coordinates were saved every 100 ps for analysis. The simulations were conducted in an isothermal and isobaric nature, with the pressure maintained using an isotropic position scaling algorithm with the pressure relaxation time fixed at 2 ps. Long-range electrostatics were calculated by a particle mesh Ewald method with a 10 Å cut-off {Darden, 1993 #5557}. Three replica simulations for each of the systems were performed. The results of the MD simulations were analyzed using various tools and methods, including the built-in utility of the GROMACS program from Groningen University, and in-house scripts.

### Binding free energy calculations

The MM-GBSA module, which is implemented in Amber18, was performed to calculate the binding free energy of PIP2 with the Kir2.1 channel and its mutants. The binding free energy can be decomposed into contributions from individual interacting residues of the channels, which consist of four energy terms: non-bonded electrostatic interaction, van der Waals energy in gas phase, polar, and non-polar solvation free energies. The polar component is calculated using a Generalized Born (GB) implicit solvation model. The non-polar component is calculated using a solvent accessible surface area model. 100 snapshots were extracted from the 90-100ns MD simulation trajectory at an interval of 100ps for the binding free energy calculations.

### Quantification and Statistical Analysis

Data were analyzed using Clampfit, GraphPad (Prism), and Excel software. Quantification and analysis approaches were specific in each experiment and are described in the Figure legends. Data were assessed for statistical differences between groups by unpaired Mann-Whitney rank test or two-tailed Students t-test, as indicated in the Figure legends, following interrogation of variance with Bonferroni post hoc analysis to test differences within pairs of group means for all datasets with an F-value of p < 0.05. Data are presented, where indicated as the mean ± standard deviation (s.d.). The number of replicates for each study are described in the legends.

## Results

### Hypoxic inhibition of cardiac *I_K1_* is precluded by TAK-981

To measure the effect of hypoxia on *I_K1_,* we recorded whole-cell currents from rat ventricular cardiomyocytes (RVCMs) and reduced the O_2_-tension of the extracellular superfusate from ambient levels (∼21%) to 2% O_2_. Consistent with our prior studies, acute hypoxia evoked a rapid decrease in *I_K1_* to 57 ± 3% of control levels (mean ± SEM, n = 10, P<0.0001, paired t-test) and the residual current was blocked by 3 mM Ba^2+^ (**Figure, 1A**) ^15^. We have previously shown that hypoxic inhibition of *I_K1_* is caused by rapid SUMOylation of Kir2.1 channels, the primary contributors to inward K^+^ current in ventricular cardiomyocytes ^15^. To test whether this effect could be blocked by SUMO pathway modulators, we pretreated RVCMs with TAK-981, a potent, small-molecule inhibitor of subunit-1 of the SUMO E1-activating enzyme, SAE1 ^32^. A 3-hour pretreatment with 100 nM TAK-981 increased *I_K1_* by ∼26% from 176 ±12 pA/pF to 222 ± 11 pA/pF (n = 10, P<0.01, unpaired t-test). These data suggest that *I_K1_* is suppressed by basal SUMOylation, and that this inhibition is relieved when the SUMO-pathway is blocked. Furthermore, the TAK-981 treatment made *I_K1_* insensitive to acute hypoxia, indicating that active SAE1 is necessary for the hypoxic regulation of the current in cardiomyocytes (**Figure, 1A**) ^15^.

**Fig 1.**
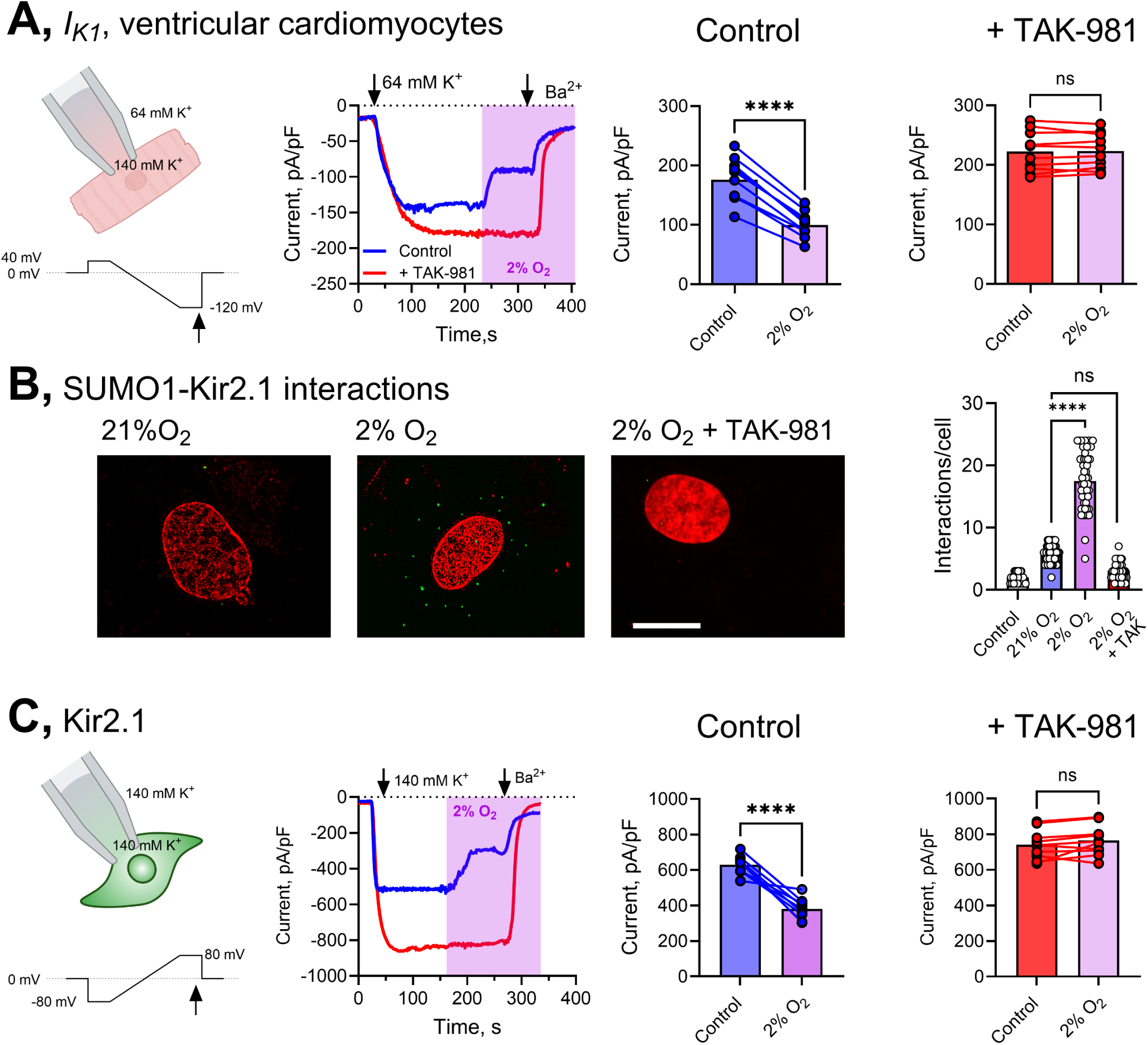
Hypoxic inhibition of cardiac I_K1_ is precluded by TAK981. Patch-clamp data are from 10-12 cells per group. *I_K1_* was evoked in rat ventricular cardiomyocytes (RVCMs) using a high external K^+^ recording buffer and measured at -120mV. Kir2.1 currents from heterologous HEK293T cells were measured at -80 mV. Hypoxia is a drop in O_2_ from ambient levels to 2% (pink box), measured at the cell. TAK-981 was used at 100 nM and was applied 3 hours before the experiment. Statistical differences were determined using a paired, two-tailed Student’s t test where **** is p< 0.0001. *See also Figures S1, S2, S3, and S4* **A,** *Left*, Cartoon representation of the method to study *I_K1_*, with representative time-courses showing hypoxic-inhibition of *I_K1_* (blue). Pretreatment TAK-981 increases basal *I_K1_* and precludes the effects of hypoxia (red). *Right*, Summary histograms showing hypoxic inhibition of *I_K1_* for control and TAK-981 treated RVCMs. **B,** Representative confocal fluorescence micrographs of RVCMs showing proximity ligation reaction between Kir2.1 and SUMO1 in green and the nucleus is red. Summary histogram for at least 20 cells per condition. Control is no primary antibody. **C,** *Left*, Cartoon representation of the method to Kir2.1 channels expressed in HEK293T cells, with representative time-courses showing hypoxic-inhibition of the current (blue). The residual current is blocked by 3 mM Ba^2+^. Pretreatment with TAK-981 increases basal Kir2.1 current and precludes the effects of hypoxia (red). *Right*, Summary histograms showing hypoxic inhibition of Kir2.1 for control and TAK-981 treated cells.

### TAK-981 prevents hypoxia-induced SUMOylation of Kir2.1 channels

Kir2.1 passes the largest fraction of *I_K1_* in RVCMs and is the only component of the current that is sensitive to SUMOylation ^15^. To determine if TAK-981 inhibits Kir2.1-SUMOylation we visualized and quantified the interactions between native SUMO1 and Kir2.1 channels in RVCMs using a proximity-ligation association (PLA) assay, based on antibodies that we validated previously ^15^. Under ambient levels of O_2_, PLA detected an average of 6 ± 1 interactions between Kir2.1 and SUMO1 in RVCMs. Exposing the cells to hypoxia for 2-mins increased PLA interactions by 3-fold to 18 ± 4 (P<0.001) per cell, indicating rapid SUMOylation of Kir2.1. In contrast, pretreatment with TAK-981 reduced the basal PLA-interaction rate to background levels (fewer than 3-interactions per cell) and blocked the hypoxia-induced increase in Kir2.1-SUMOylation (**Figure, 1B**).

To measure the effect of TAK-981 on Kir2.1 function, we expressed the channels in HEK293T cells. First, we showed that an acute application of 100 nM TAK-981 did not alter the activity of Kir2.1 channels. Similar results were obtained using Kir2.1 channels expressed in *Xenopus laevis* oocytes (**Figure S1A-B**). Consistent with our findings in cardiomyocytes, hypoxia evoked a ∼40% reduction in Kir2.1 currents in HEK293T cells (n = 10, P<0.0001, paired t-test). However, in cells pretreated with TAK-981, Kir2.1 currents were increased by ∼30% from 628 ± 18 pA/pF to 821 pA/pF (n = 10, P<0.0003, unpaired t-test), and were insensitive to hypoxic regulation (**Figure, 1C**).

We previously demonstrated that SUMOs modify Kir2.1 at residue K49 on the cytoplasmic face of the channel. Therefore, K49 substitutions prevent SUMOylation of Kir2.1 ^15^. As expected, Kir2.1-K49Q channels passed currents ∼35% larger than those recorded from wild type channels. Additionally, the Kir2.1-K49Q currents were insensitive to acute hypoxia and were not altered when the cells were pretreated with TAK-981 (**Figure S1C**).

To study the association between Kir2.1 and SUMO1 in live cells, we measured donor-decay Förster Resonance Energy Transfer (FRET) between mTFP1-tagged Kir2.1 and YFP-SUMO1. mTFP1-Kir2.1 localized to the plasma membrane and photobleached under continual illumination, with a time constant (tau) of 54 ± 1 seconds, in the presence of free YFP. The tau increased by 68.5% to 92 ± 2 s when mTFP1-Kir2.1 was studied in the presence of YFP-SUMO1, indicating SUMOylation. A similar result was obtained with YFP-SUMO2. However, pretreatment with TAK-981 abolished FRET between mTFP1-Kir2.1 and YFP-tagged SUMO1 or SUMO2, consistent with inhibition of SAE1. In contrast, robust FRET between mTFP1-Kir2.1 and the SUMO E2 conjugase, Ubc9, remained unaffected by TAK- 981 pretreatment (**Figure, S1D-E**). FRET was not observed between mTFP1-tagged Kir2.1-K49Q and YFP-tagged SUMO1 or SUMO2. However, the K49Q mutation did not interfere with the interaction between the channel and YFP-Ubc9, indicating that the SUMOylation motif is not required for Ubc9 to associate with Kir2.1 (**Figure, S1F**).

### The Ubc9 inhibitor 2-D08 opposes hypoxia-induced SUMO regulation of Kir2.1 channels

To confirm that hypoxic-induced SUMOylation could be blocked by inhibitors that target components of the SUMO-pathway downstream of the E1-activating enzyme, we also studied 2’,3’,4’- trihydroxyflavone (2-D08), which inhibits the SUMO-specific E2 conjugase, Ubc9 with an IC_50_ of 6 μM ^33, 34^. Like TAK-981, pretreatment with 30 µM 2-D08 increased Kir2.1 channel currents and made them insensitive to hypoxia (**Figure, S2A-B**). Cells treated with 2-D08 also showed no donor-decay FRET between mTFP1-Kir2.1 and YFP-tagged SUMO1 or SUMO2, but FRET interactions with Ubc9 or the catalytically inactive Ubc9-C93S were preserved. Notably, the interaction between Ubc9 and Kir2.1- K49Q remained intact in the presence of 2-D08 (**Figure, S2C-D**).

### TAK-981 pretreatment prevents acute SUMO-regulation of Kir2.1

Next, we examined whether TAK-981 could prevent the inhibition of Kir2.1 under two additional conditions known to promote SUMOylation of ion channels. First, we introduced purified SUMO1_97_ protein into cells via the recording pipette ^11, 15, 35–37^. Since SUMO1_97_ retains the C-terminal diglycine motif, it can interact with endogenous SUMO-pathway enzymes, including SAE1. As observed in previous studies, 1 µM SUMO1_97_ reduced Kir2.1 current amplitude by ∼45%, leaving a residual current that was insensitive to hypoxia (**Figure, S3A**). However, when cells were pretreated with TAK- 981, SUMO1_97_ no longer reduced Kir2.1 currents (**Figure, S3B**), indicating that purified SUMO1_97_ proteins require SAE1 to regulate Kir2.1 channels.

Next, we tested whether TAK-981 could counteract the effects of N106, a small molecule activator of SAE1 that promotes SUMOylation ^38^. Pretreatment with 10 µM N106 reduced Kir2.1 currents by ∼40% compared to untreated cells and abolished hypoxic inhibition. Including 1 µM SUMO1_97_ into N106-treated cells further suppressed Kir2.1 currents, from 375 ± 13 pA/pF to 200 ± 17 pA/pF (n = 9, P<0.001, unpaired t test). These effects were blocked by pretreatment with TAK-981 (**Figure, S3C-F**). Furthermore, TAK-981 prevented FRET interactions between mTFP1-tagged Kir2.1 and YFP-tagged SUMOs, but not YFP-Ubc9, in N106 treated cells (**Figure, S3G**). As expected, N106 did not affect the current magnitude or FRET interactions of Kir2.1-K49Q variant channels (**Figure, S4**).

### Kir2.1 channels can carry up to two SUMO1 monomers

We used total internal reflection fluorescence (TIRF) microscopy to measure the number of SUMO1 monomers associated with individual Kir2.1 channels. First, we observed that mTFP-Kir2.1 (mT- Kir2.1) and mCherry-SUMO1 (mCh-SU1) colocalize at the plasma membrane of live HEK293T cells (**Figure, 2A**). Under continual TIRF illumination, fluorophores bleach in a stepwise manner that corresponds to their stoichiometry. For colocalized particles, mTFP-Kir2.1 bleached in 4-steps, while mCherry-SUMO1 bleached in two, indicating a maximum stoichiometry of two SUMO1 monomers per channel. In contrast, mCherry-SUMO1 did not colocalize with mTFP-Kir2.1-K49Q (mT-K49Q) channels (**Figure, 2A**).

**Fig 2.**
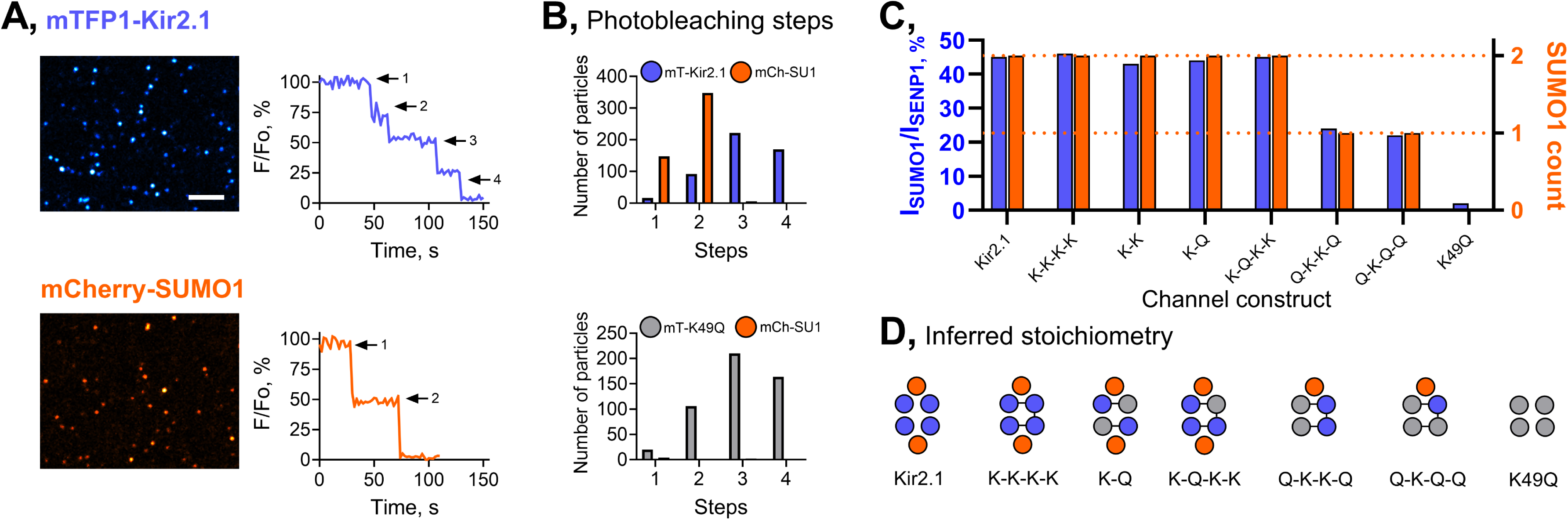
Kir2.1 channels are modified by up to two SUMO1 monomers. mTFP1-tagged Kir2.1 (cyan) and mCherry-SUMO1 (orange) were expressed in HEK293T cells and visualized at the membrane by TIRF microscopy. The stoichiometry of single Kir2.1-SUMO1 complexes was measured by single particle photobleaching. K49Q is the mutation in Kir2.1 that prevents SUMO conjugation. *See also Figure S5* **A,** Representative TIRF micrographs and fluorescence photobleaching records showing that single particles containing mTFP1-Kir2.1 photobleaching in 4-steps (*upper*, cyan) and mCherry-SUMO1 photobleaching in 2 steps (*lower*, orange). **B,** *Upper*, Summary histogram showing the 4 to 2 stoichiometry of colocalized particles with mTFP- Kir2.1 (mT-Kir2.1) and mCherry-SUMO1 (mCh-SU1). *Lower*, Summary histogram for mTFP-Kir2.1- K49Q (gray) showing that the channel does not localize with mCherry-SUMO1. **C,** Summary histogram showing the relative SUMO-sensitive current for Kir2.1 concatemers (K represents wild type subunits, Q is K49Q), correlated with the number of mCherry-SUMO1 monomers associated with the channel. **D,** Cartoon representations of the inferred stoichiometries of Kir2.1 concatemers. Wild type Kir2.1 subunits (K) are in blue, K49Q subunits (Q) in gray, and SUMO1 is in orange.

To determine the relative position of SUMO1 monomers on the channels, we utilized concatenated Kir2.1 subunits to control the location of SUMOylation motifs within the tetramer. Channels formed by dimers or linked tetramers of wild type subunits passed similar levels of current to Kir2.1 monomers (Figure, 2C). Co-expression of SUMO1 and Ubc9 reduced the current by ∼45% compared to expression of the channel with the deSUMOylating enzyme SENP1, which suppresses background SUMOylation of ion channels, including Kir2.1 ^11, 15, 35, 39^. TIRF photobleaching confirmed the presence of two SUMO1-monomers per channel (**Figure, 2C**; **Figure S5**).

Next, we introduced Kir2.1-K49Q subunits into these constructs to selectively eliminate SUMO-binding sites within the channel tetramers. First, we studied cells expressing dimers formed by linking Kir2.1 wild type (WT) subunits to Kir2.1-K49Q subunits (WT-K49Q). Channels formed by these dimers passed currents with a similar amplitude to currents from wild type (WT-WT) dimers. Both WT-WT and WT-K49Q dimers, tagged with one mTFP1 fluor per dimer, formed channels that photobleached in two steps. When SUMO1 was co-expressed, currents from WT-WT and WT-K49Q dimers were suppressed by ∼45%, with two SUMO1 monomers associated with each channel (**Figure, 2C-D**; **Figure S5**).

We then studied tetrameric constructs with three, two or one SUMO1 binding sites. TIRF photobleaching showed that channels with three SUMO binding sites associated with two SUMO1 monomers when the available sites are adjacent, suggesting that Kir2.1 channels are SUMOylated on diagonally opposite subunits. As expected, channels with one SUMOylation site carried a single SUMO1. These results also confirm that mCherry-tagged SUMO1 does not form chains in this system. Functional studies using the same concatenated constructs revealed that SUMOylation by a single SUMO1 monomer inhibits ∼22% of maximal Kir2.1 current (**Figure, 2C-D**; **Figure S5)**.

### TAK-981 and SENP1 partially recover the Kir2.1-R67Q current phenotype

ATS1 is a rare multisystem channelopathy associated with mutations in *KCNJ2*, the gene encoding for Kir2.1 ^6^. Structural analysis of the Kir2.1 channel revealed that the SUMO-target lysine, K49, is located near the ATS1 disease locus, R67 (**Figure, 3A**). Given these observations, we hypothesized that SUMOylation could augment the R67Q channel phenotype.

In common with previous studies, we found that homotetrameric Kir2.1-R67Q or Kir2.1-R67A channels with a homozygous genotype, are non-functional (**Figure, S6A-B**). However, ATS1 is typically heterozygous, with an autosomal dominant, loss-of-function phenotype. To reflect this, we studied channels formed from dimers with a wild type Kir2.1 subunit linked to a Kir2.1-R67Q subunit (WT-R67Q), which pass measurable whole-cell currents ^1, 18, 40, 41^. Of note, channels formed by WT-WT dimers behaved similarly to those formed by WT monomers, passing a mean current of 634 ± 15 pA/pF (**Figure, S6D**). In contrast, currents from channels formed by WT-R67Q dimers were ∼85% smaller and decreased by an additional ∼65% from 212 ± 5 pA/pF to 89 ± 4 pA/pF when exposed to hypoxia (n = 8, P < 0.0001, paired t-test; **Figure, 3B-C**). However, WT-R67Q currents increased by ∼50% when cells were pretreated with TAK-981 or when 2 µM SENP1 was included in the recording pipette, and these currents were resistant to hypoxia (**Figure, 3C, E**).

Donor-decay FRET photobleaching studies confirmed that mTFP1-tagged Kir2.1-R67Q channels interacted with YFP-tagged SUMO1 and SUMO2 in control cells, but not in TAK-981-treated cells. Kir2.1-R67Q also interacted with YFP-Ubc9, and this interaction was unaffected by TAK-981 (**Figure S6C**).

Since SUMOylation can occur at WT or R67Q subunits, we generated K49Q-R67Q dimers to localize SUMOylation to the subunit carrying the ATS1 mutation. Currents measured from channels formed by K49Q-R67Q dimers were ∼30% smaller than those of WT-R67Q channels and showed a further ∼50% reduction under acute hypoxia, from 162 ± 3 pA/pF to 78 ± 3 pA/pF (n = 12, P < 0.001, paired t-test). Pretreatment with TAK-981 significantly increased K49Q-R67Q channel currents by ∼93% to 314 ± 4 pA/pF (n = 12, P < 0.0001, unpaired t-test) and blocked the effects of hypoxia. Similar results were observed when K49Q-R67Q channels were studied with 2 µM SENP1 in the recording pipette (**Figure 3D, E**).

**Fig 3.**
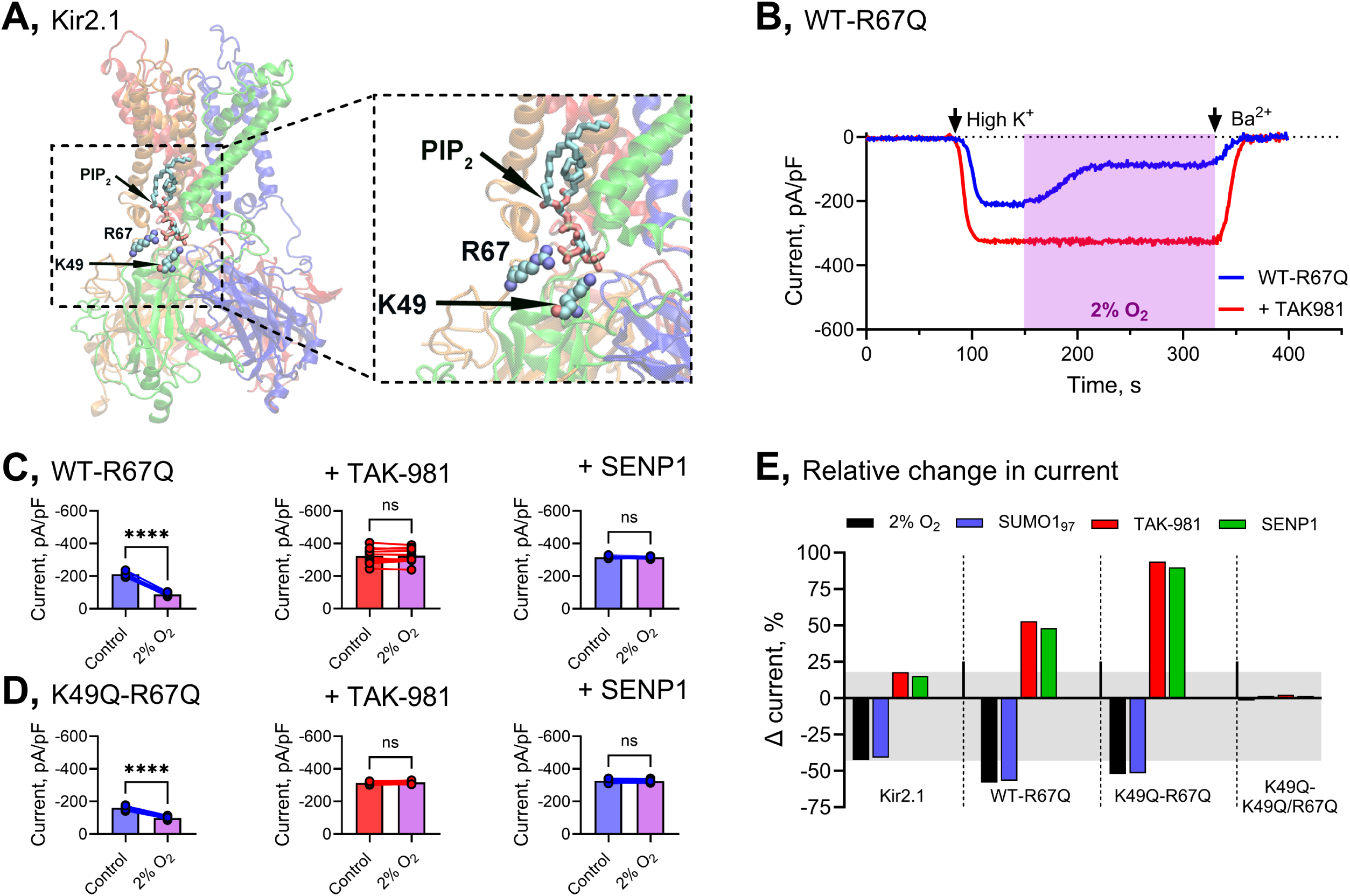
Inhibiting SUMOylation increases Kir2.1-R67Q channel currents. Patch-clamp data are from 10-12 cells per group. Currents from channels formed by Kir2.1-dimers were measured at -80 mV, per the Methods. Hypoxia is a drop in O_2_ from ambient levels to 2% (pink box), measured at the cell. TAK-981 was used at 100 nM and was applied 3 hours before the experiment. Purified SUMO1_97_ was used at 1 µM and SENP1 was used at 2µM and was included in the recording pipette where indicated. Statistical differences were determined using a paired, two-tailed Student’s t test where **** is p< 0.0001. *See also Figure S6* **A,** Molecular model of Kir2.1 based on PDB 7ZDZ with docked PIP_2_ showing the position of K49 and R67 on the same subunit of the channel. **B,** Representative time-courses showing hypoxic-inhibition of channels formed from a wild type Kir2.1 (WT) and R67Q-subunit dimer (WT-R67Q; blue). The residual current is blocked by 3 mM Ba^2+^. Pretreatment with TAK-981 increased the Kir2.1 current magnitude and precluded the effects of hypoxia (red). **C,** Summary histograms showing that hypoxic inhibition of WT-R67Q channels is precluded by TAK- 981 or SENP1. **D,** Summary histograms showing that hypoxic inhibition of K49Q-R67Q channels is precluded by TAK-981 or SENP1. **E,** Summary chart showing the relative change in current for various Kir2.1 constructs in response to hypoxia (black), purified SUMO1_97_ (blue), TAK-981, or purified SENP1 (red). The shaded area indicates the response range of wild type Kir2.1 channels.

SUMO-insensitive K49Q-K49Q/R67Q dimers produced currents that are ∼50% the magnitude of those measured from wild type Kir2.1 channels and were resistant to modulation by hypoxia, SUMO1_97_, TAK-981, or SENP1 (**Figure S6F**).

### PIP_2_ interactions in ATS1-variant Kir2.1 channels

Kir2.1 channel gating requires interactions with PIP_2_ ^17, 19^. Several ATS1 mutations, including R67Q, are proposed to impair Kir2.1 currents by disrupting the channel’s interface with PIP_2_ ^1, 18, 41, 42^. Kir2.1- PIP_2_ interactions are also disrupted by SUMOylation ^15^, leading us to investigate whether SUMOylation exacerbates the R67Q current phenotype by further destabilizing PIP_2_ regulation of the channel.

To interrogate the mechanistic basis for the impact of the R67Q mutation and SUMOylation on Kir2.1, we modeled multiple configurations of the channel to determine changes to the PIP_2_ binding pose. In the wild-type Kir2.1 channels, the phosphate headgroups of PIP_2_ interact with multiple residues between adjacent subunits, including K49 and R67 (**Figure 4A-C**). In channels formed by WT-R67Q dimers, interactions with K49, Q67 and K185 are predicted to be lost, and interactions with R80 and W81 are predicted to be weakened. However, new interactions are predicted to be established at R82 and R218. These changes reduce the overall binding free energies for PIP_2_ from -91.94 kcal/mol in the wildtype system to -79.73 kcal/mol in the WT-R67Q system (**Figure 4A-C**).

**Fig 4.**
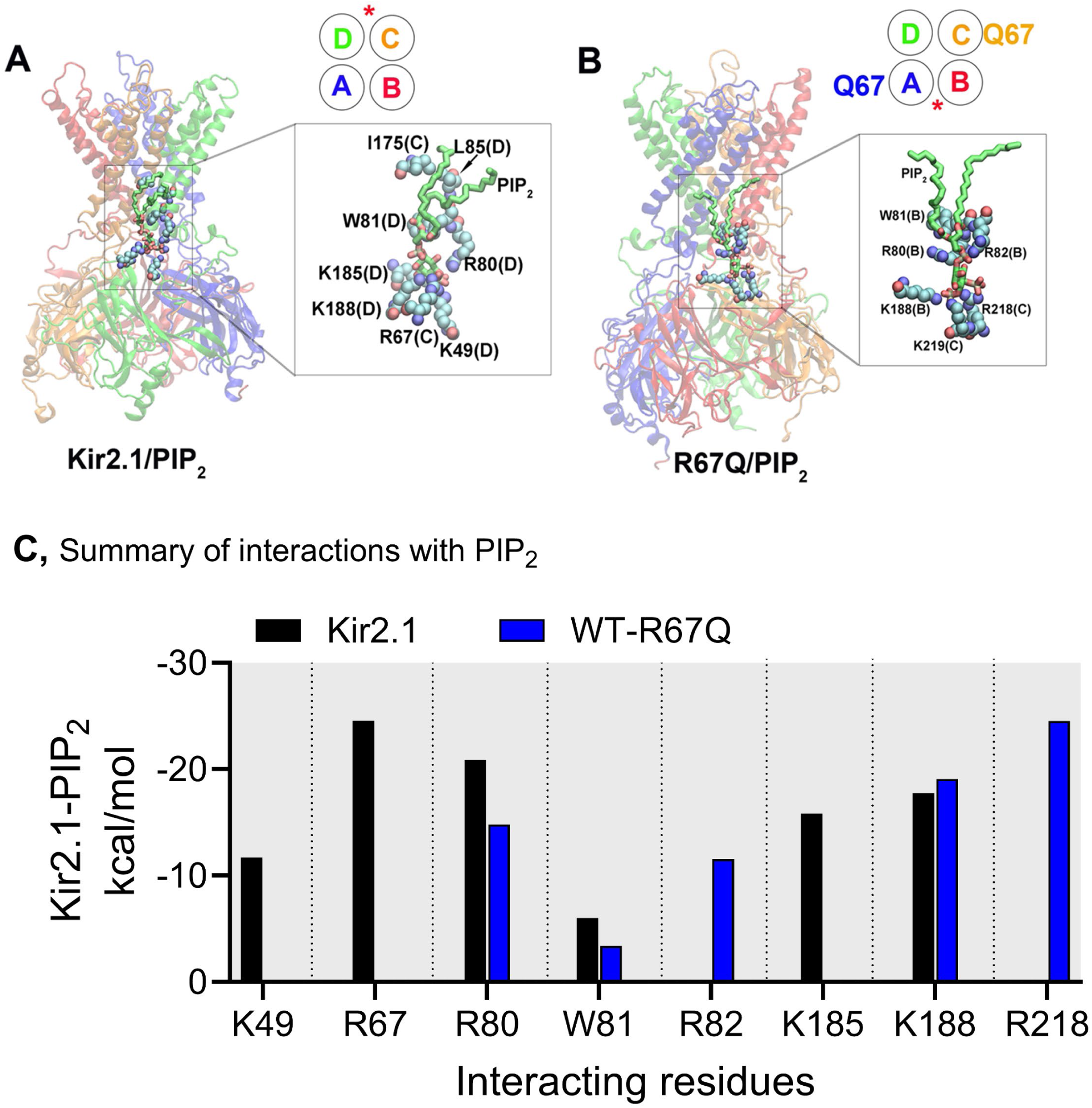
The energetics of the PIP_2_ interaction interface. Molecular models of Kir2.1 with PIP_2_ were generated by MD simulations. The energy of interaction between PIP_2_ and its coordinating residues was calculated using the MM-GBSA module of Amber program. **A,** PIP_2_ binding pose in the Kir2.1 channel. The inset shows one PIP_2_ between subunits C and D. **B,** PIP_2_ binding pose in Kir2.1 with the R67Q mutations in subunits A and C. The inset shows one PIP_2_ between subunits B and C. **C,** Summary histogram showing the energy of interaction between PIP_2_ and the residues indicated for the two channel systems: wild type Kir2.1 (black) and alternating wild-type, R67Q subunits (WT- R67Q, blue).

### SUMOylation enhances disruption of Kir2.1-R67Q interactions with PIP_2_

To measure the functional effects of the R67Q mutation, hypoxia, and direct-SUMOylation on Kir2.1- PIP_2_ interactions, we used optogenetic activation of an inositol 5-phosphatase to dephosphorylate PIP_2_ in voltage-clamped cells ^43, 44^. Cells were co-transfected with the channel, CRY2-5-ptase_OCRL_ (5- ptase), and its cognate the membrane anchor, CIBN-CAAX. Photoactivation of the system using 460 nm light rapidly recruited 5-ptase to CIBN, leading to 5-dephosphorylation of PIP_2_ and production of PI(4)P, a phospholipid that does not support efficient Kir2.1 gating (**Figure 5A**) ^45^. Therefore, photoactivation of 5-ptase caused a rapid mono-exponential reduction in current from ∼625 to 245 pA/pF for Kir2.1 channels formed from wild type monomers or link dimers (WT-WT), with a time constant (τ_ptase_) of ∼36 seconds (**Figure 5B-D**). Pretreatment with TAK-981 prolonged τ_ptase_ to ∼43 seconds for both wild type Kir2.1 channel constructs. In contrast, τ_ptase_ accelerated to ∼16 seconds under acute hypoxia, with SUMO1_97_, or both. These effects were blocked by TAK-981 pretreatment (**Figure 5C-D**).

**Fig 5.**
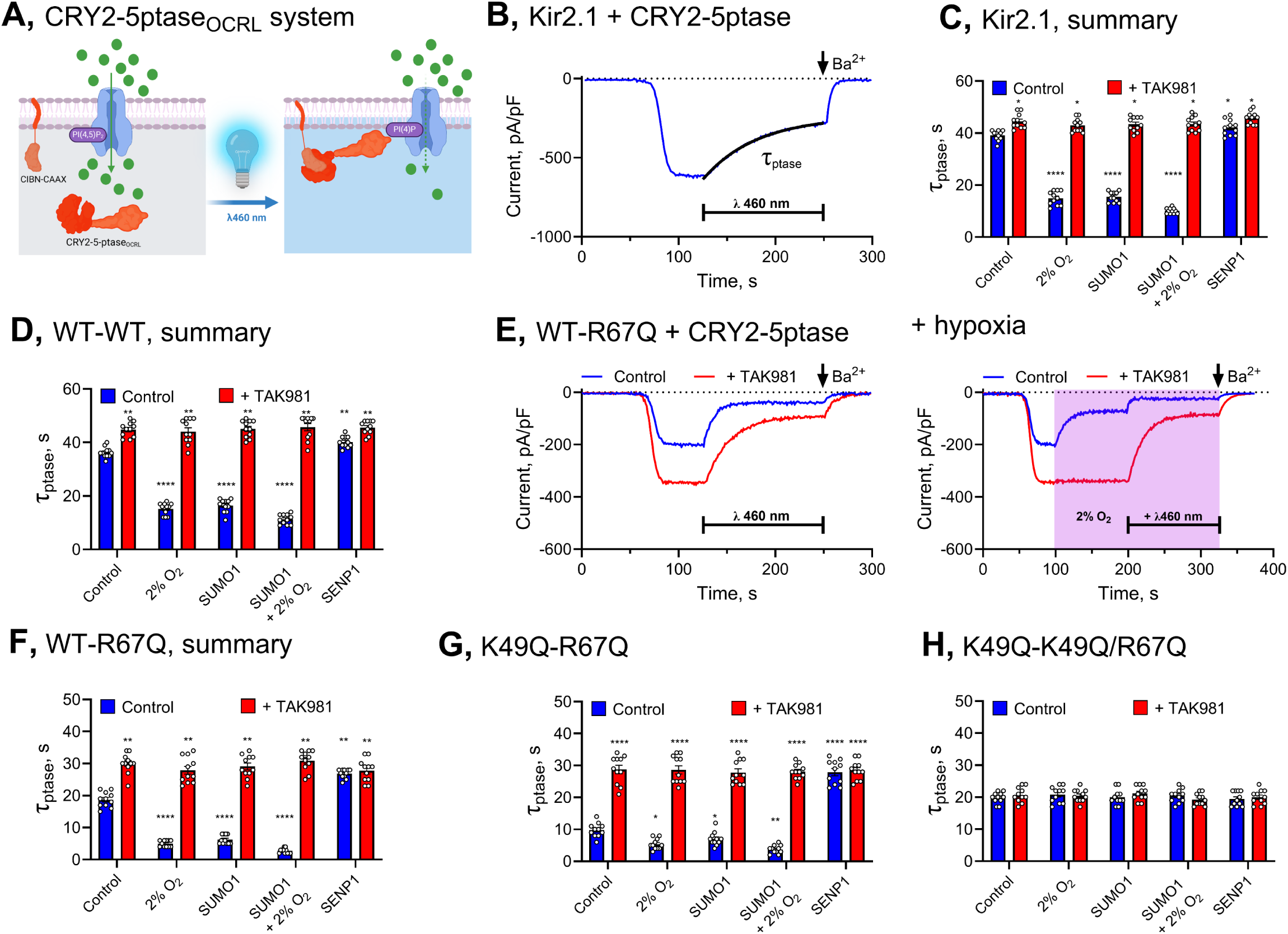
SUMOylation enhances the disruption of Kir2.1-R67Q - PIP_2_ interactions. Kir2.1 channel currents were measured from HEK293T cells by whole-cell patch clamp recording with simultaneous optogenetic dephosphorylation of PIP_2_. Twelve-15 cells were studied per condition. Hypoxia is a drop in O_2_ from ambient levels to 2% (pink box), measured at the cell. TAK-981 was used at 100 nM and was applied 3 hours before the experiment. Purified SUMO1_97_ was used at 1 µM and SENP1 was used at 2 µM and was included in the recording pipette where indicated. **A,** Cartoon showing the optogenetic platform: Cells are transfected to express the 5-phospholipid phosphatase, OCRL fused to CRY2 (CRY2-5ptase) and the CIBN, a protein partner for CRY2, anchored to the membrane via a CAAX motif. In the presence of 460 nm (blue) light, CRY2 fuses to CIBN rapidly, bringingCRY2-5ptase to the inner leaflet of the plasma membrane. **B,** Representative time-course showing the monoexponential fit to the decrease in Kir2.1 channel current evoked by photoactivation of CRY2-5ptase with time constant τ_ptase_. **C,** Summary histograms showing τ_ptase_ determined for Kir2.1 channels studied under the conditions indicated (blue) and following pretreatment with TAK-981 (red). **D,** Summary histograms showing τ_ptase_ determined for Kir2.1 channels formed by WT-WT dimers, studied under the conditions indicated (blue) and following pretreatment with TAK-981 (red). **E,** *Left*, Representative time-courses for currents measured from WT-R67Q dimers showing the τ_ptase_ determined from control (blue) cells and from TAK-981 pretreated cells (red). *Right*, The experiment is repeated with cells exposed to hypoxia. **F,** Summary histograms showing τ_ptase_ determined for channels formed by WT-R67 dimers, studied under the conditions indicated (blue) and following pretreatment with TAK-981 (red). **G,** Summary histograms showing τ_ptase_ determined for channels formed by K49Q-R67 dimers, studied under the conditions indicated (blue) and following pretreatment with TAK-981 (red). **H,** Summary histograms showing τ_ptase_ determined for channels formed by K49-K49Q/R67 dimers, studied under the conditions indicated (blue) and following pretreatment with TAK-981 (red).

For WT-R67Q channels, τ_ptase_ was more rapid (16 ± 2 seconds, n = 11), consistent with decreased stability of channel-PIP_2_ interactions. The τ_ptase_ doubled to 30 ± 2 seconds (n = 11, P < 0.001, unpaired t-test) with TAK-981 pretreatment, suggesting stabilization of channel-PIP_2_ interactions (**Figure 5E- F**). Hypoxia significantly accelerated τ_ptase_ for WT-R67Q to 5 ± 3 seconds (n = 11), as did SUMO1_97_ (6 ± 2 seconds). When hypoxia and SUMO1_97_ were combined, τ_ptase_ was reduced further to 3 ± 1 seconds. These effects were abolished by TAK-981 (**Figure 5E-F**). Similarly, including 2 µM SENP1 in the recording pipette prolonged τ_ptase_ to 27 ± 2 seconds (n = 11), with no further change when TAK-981 was applied (**Figure 5F**).

We also measured τ_ptase_ values from cells expressing K49Q-R67Q channels, which restrict SUMOylation to the R67Q-containing subunit. Notably, τ_ptase_ for K49Q-R67Q was accelerated to 10 ± 2 seconds (n = 11), indicating further destabilization of the channel-PIP_2_ interface. Hypoxia accelerated τ_ptase_ to 5 ± 3 seconds, as did SUMO1_97_ (n = 11). These effects were blocked by TAK-981 or SENP1 (**Figure 5G**). When both subunits of the dimer were insensitive to SUMOylation (K49Q- K49Q/R67Q), τ_ptase_ was 32 ± 3 seconds (n = 9) and was unaffected by hypoxia, SUMO1_97_, TAK-981, or SENP1 (**Figure 5H**).

## Discussion

This study identifies a synergistic mechanism by which the ATS1-linked Kir2.1-R67Q mutation and hypoxia converge to suppress Kir2.1 function and reduce *I_K1_*. The resulting suppression of Kir2.1 supports a two-hit hypothesis in ATS1, in which a genetic predisposition is amplified by a stress-induced post-translational modification. Mechanistically, we demonstrate that both hypoxia and the R67Q mutation impair Kir2.1-PIP_2_ interactions, a gating requirement for the channel. Hypoxia exerts this effect through rapid SUMOylation of Kir2.1 at lysine 49, while R67Q reduces channel–PIP₂ affinity.

Inhibition of the SUMO pathway with TAK-981 reverses these effects, reduces basal SUMOylation, and restores current through wild-type and R67Q channels.

Most pathogenic *KCNJ2* mutations associated with ATS1 are proposed to disrupt Kir2.1–PIP₂ interactions ^1, 4, 18^. Our findings suggest that environmental stressors, including hypoxia, exacerbate this loss-of-function phenotype via a shared SUMOylation-dependent mechanism. This raises the possibility that other arrhythmogenic stimuli such as exposure to carbon monoxide or the gasotransmitter hydrogen sulfide, that act to disrupt Kir2.1-PIP_2_ interactions might also expand the inherited risk of an ATS1 channelopathy via a similar two-hit mechanism ^19, 46^.

Kir.2.1-PIP_2_ binding is governed by electrostatic interactions between negatively charged PIP_2_ (-5e) and conserved basic residues at four binding pockets within the channel. The R67Q mutation weakens these interactions, as confirmed by our binding free energy calculations, and this destabilization is magnified when SUMOylation is constrained to the same subunit, as in our concatemeric constructs. While Kir2.1 exhibits low basal SUMOylation under normoxia ^15^,, its suppressive impact is significantly enhanced in the R67Q background, leading to a disproportionate reduction in current. Accordingly, R67Q-containing channels show a marked increase in current under deSUMOylating conditions, either by SUMO-pathway inhibition or SENP exposure.

Although each Kir2.1 subunit contains a conserved K49 SUMOylation site, our findings indicate that only two SUMO1 monomers are accommodated per channel, positioned on diagonally opposite subunits. This stoichiometry, consistent with our prior work on Kv2.1 and Kv7.1, likely reflects steric constraints on the SUMOylation machinery. Functionally, this results in a graded suppression of current: ∼20% with a single SUMO-modified subunit and up to ∼40% with two. In the R67Q background, this suppression is further amplified, consistent with our observation that hypoxia has a more severe inhibitory effect on mutant channels ^11, 14, 15, 39, 48, 49^.

We also show that SUMOylation is dynamically regulated by cellular stress. In addition to hypoxia, SUMOylation is potentiated by exposure to SAE1 activators, purified SUMO1 protein, and adverse conditions such as oxidative stress, inflammation, and heat shock ^50–52^. The molecular mechanisms that mediate these effects remain incompletely understood but may involve increased activity of SUMO-conjugating enzymes or reduced deSUMOylation ^20^. Given the central role of these processes in cardiovascular pathophysiology, the SUMO pathway may represent a broader integrative hub for environmental modulation of ion channel function.

Pharmacological inhibition of SUMOylation enhances *I_K1_* and mitigates hypoxia-induced current suppression. In addition to SENP1, we found that both 2-D08 and TAK-981 increase Kir2.1 currents and reverse the effects of SUMOylation^33, 34^. TAK-981, a selective and nanomolar-potent SAE1 inhibitor currently in phase 2 oncology trials ^32, 53 54^. restored Kir2.1 activity in primary ventricular cardiomyocytes and in cells expressing R67Q channels. Importantly, TAK-981 demonstrated high efficacy and selectivity, supporting its potential for repurposing in the context of arrhythmia risk reduction.

In summary, our findings demonstrate that hypoxia-induced SUMOylation and the R67Q mutation act synergistically to destabilize Kir2.1–PIP₂ interactions and suppress *I_K1_*, supporting a two-hit model of arrhythmogenesis in ATS1. Pharmacological inhibition of the SUMO pathway, particularly with TAK- 981, restores channel function and identifies Kir2.1 SUMOylation as a modifiable molecular mechanism linking metabolic stress to electrical instability in genetically predisposed patients. Future studies examining additional ATS1 variants and other SUMO-sensitive targets may further define the therapeutic potential of modulating this pathway to reduce arrhythmic risk.

## Acknowledgements

The authors thank members of the Plant, Cui and Logothetis labs at Northeastern University for constructive feedback and support. We thank the CILS core at Northeastern University for confocal microscopy support. We are grateful to Dr. Pietro De Camilli at Yale University for providing CIBN-CAAX and CRY2-5-ptase_OCRL_. The work was funded by an American Heart Association post-doctoral fellowship to K.D.G. 24POST1199293 and National Institutes of Health grant R01HL144615 to L.D.P.

## Author contributions

Electrophysiology: AC, AKY, KDG, ES FRET: AC, YX

PLA imaging: YY TIRF: YX, LDP

Constructs: TK Modeling: XM

## SUPPLEMENTAL FIGURES

**Fig S1.**
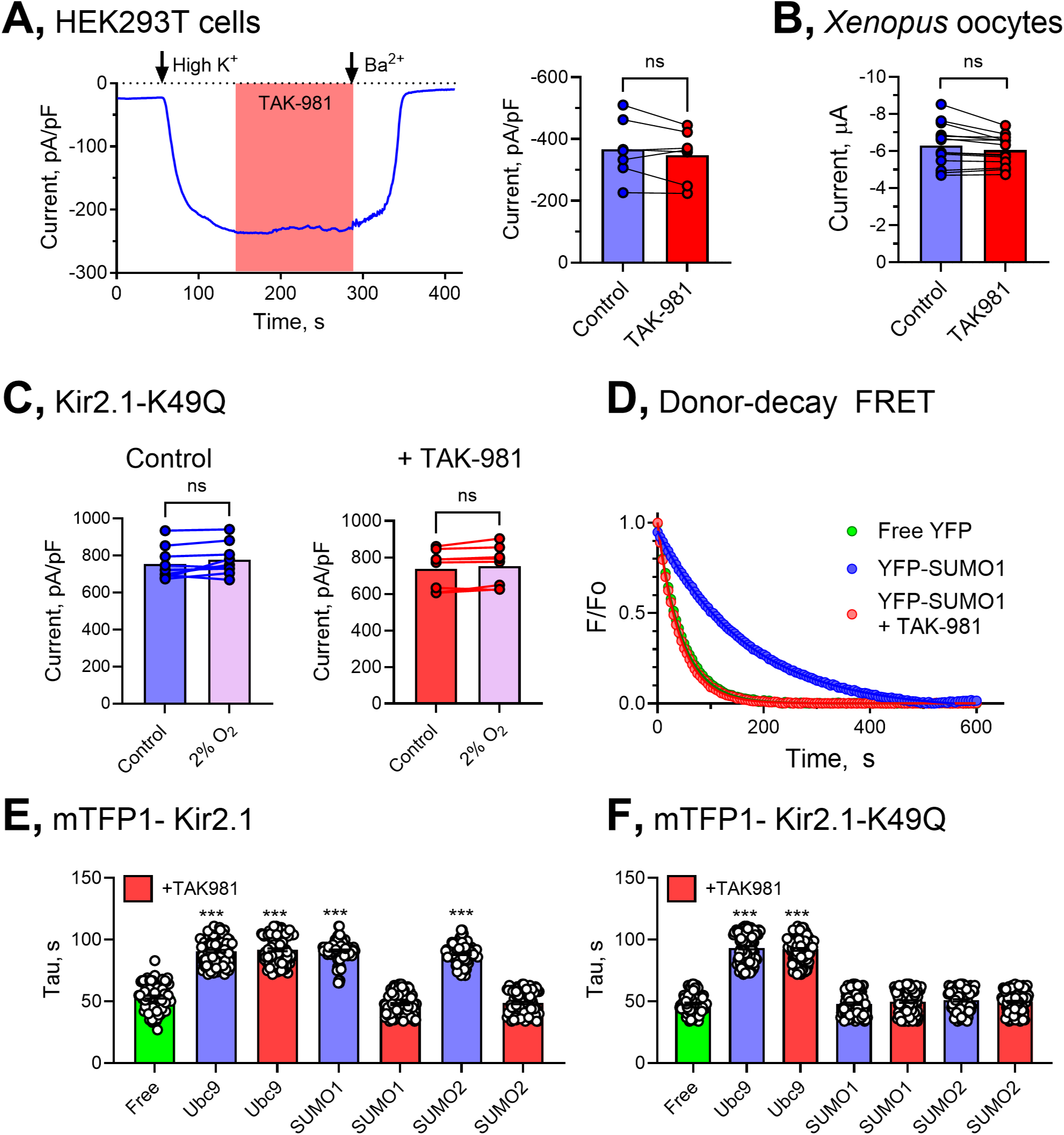
TAK-981 prevents SUMOylation of Kir2.1. FRET data are donor (mTFP) photobleaching curves, fit with a mono-exponential function to yield a fluorescence decay time constant (tau). Mean taus represent 80-120 regions of interest per group across at least 3 biological replicates. Electrophysiology data are from 6-12 cells per group. Kir2.1 currents were measured at -80 mV. Hypoxia is a drop in O_2_ from ambient levels to 2% measured at the cell. TAK-981 was used at 100 nM and was applied 3 hours before the experiment. Statistical differences were determined using a paired, two-tailed Student’s t test where *** is p< 0.001. **A,** *Left*, Representative whole-cell patch-clamp recording showing that acute application of 100 nM TAK-981 does not modulate Kir2.1 currents. *Right*, Summary histogram of paired studies showing no effect of TAK-981 on Kir2.1 current. **B,** Summary histogram of paired two-electrode voltage-clamp studies showing no effect of TAK-981 on Kir2.1 channels studied in *Xenopus* oocytes. **C,** Summary histogram showing hypoxic inhibition of SUMO-insensitive Kir2.1- K49Q channels with and without pretreatment with TAK-981. **D,** Example FRET photobleaching curves showing the decrease in mTFP1-Kir2.1 fluorescence in HEK293T cells in the presence of free YFP (green), YFP-SUMO1 (blue), or YFP-SUMO1 + TAK-981 (red). **E,** Summary histogram of photobleaching tau’s showing FRET between Kir2.1 and Ubc9, SUMO1 and SUMO2. TAK-981 (red) prevents FRET between the channel and SUMO1 and SUMO2. **F,** Summary histogram of photobleaching tau’s showing FRET between Kir2.1- K49Q but not SUMO1 or SUMO2.

**Fig S2.**
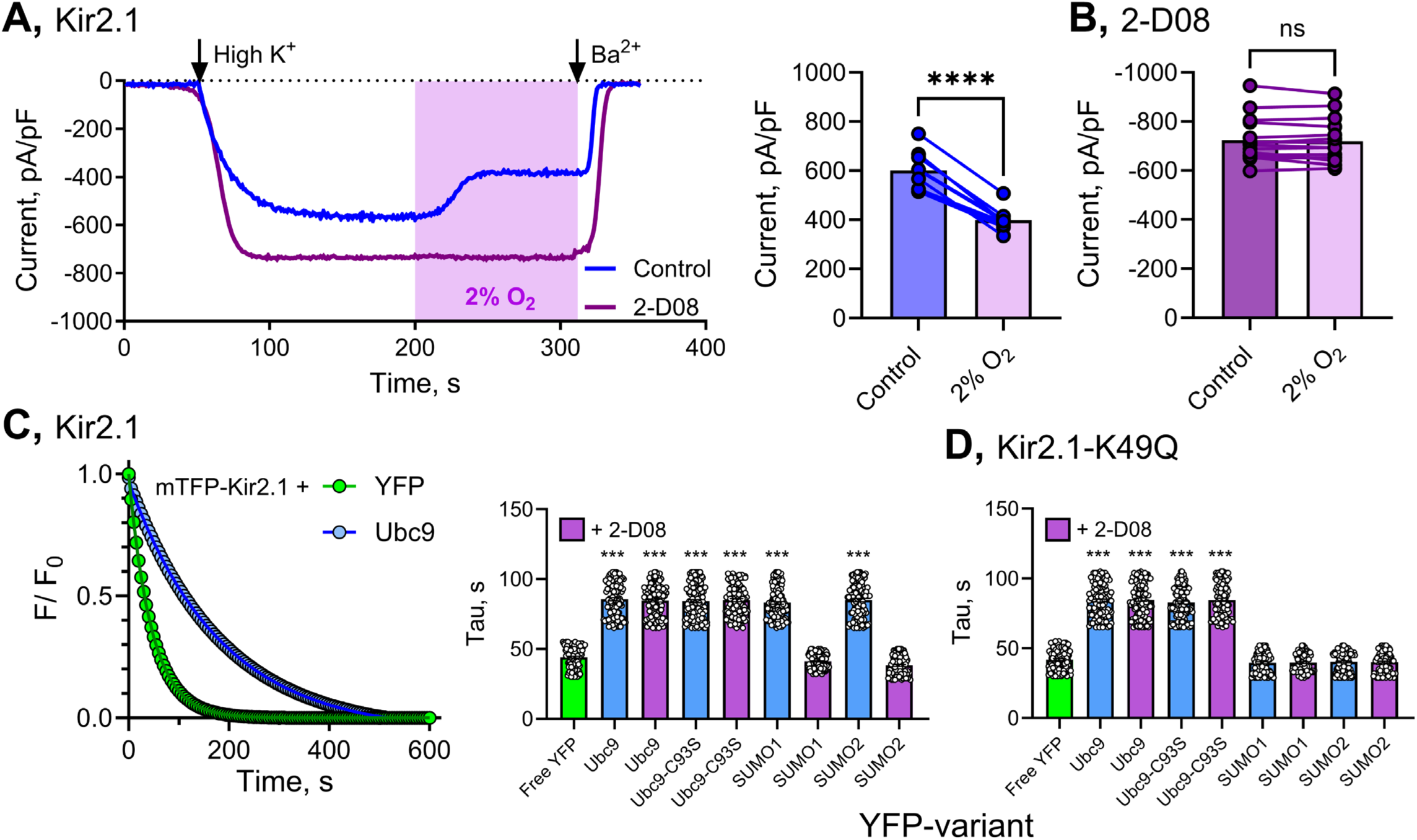
Inhibition of Ubc9 activity by 2-D08 opposes hypoxia- induced SUMO-regulation of Kir2.1 channels. FRET data aer donor (mTFP) photobleaching curves, fit with a mono-exponential function to yield a fluorescence decay time constant (tau). Mean taus represent 80-120 regions of interest per group across at least 3 biological replicates. Patch-clamp data are from 10-12 cells per group. Kir2.1 currents were measured at -80 mV. Hypoxia is a drop in O_2_ from ambient levels to 2% measured at the cell. 2-D08 was used at 30 µM and was applied 3 hours before the experiment. Statistical differences were determined using a paired, two-tailed Student’s t test where *** is p< 0.001 and **** is P<0.0001. **A,** *Left*, Representative time-courses showing hypoxic-inhibition of Kir2.1 current (blue). The residual current is blocked by 3 mM Ba^2+^. Pretreatment with 2-D08 increases basal Kir2.1 current and precludes the effects of hypoxia (purple). *Right*, Summary histogram showing hypoxic inhibition of Kir2.1 for control and **B,** 2- D08 treated cells. **C,** *Left,* Example FRET photobleaching curves showing the decrease in mTFP1-Kir2.1 fluorescence in HEK293T cells in the presence of free YFP (green) or YFP-tagged Ubc9 (blue). *Right,* Summary histogram of photobleaching tau’s showing FRET between mTFP-tagged Kir2.1 and YFP-Ubc9, or the catalytically inert variant YFP-Ubc9-C93S is not prevented by 2-D08. In contrast, FRET between the channel and YFP-tagged SUMO1 or SUMO2 is prevented by pretreatment with 2-D08. **D,** Summary histogram of photobleaching tau’s showing FRET between mTFP-tagged Kir2.1-K49Q and Ubc9, or the catalytically inert variant Ubc9-C93S is not prevented by 2-D08. In contrast, FRET is not observed between mTFP-tagged Kir2.1-K49Q and YFP- tagged SUMO1 or SUMO2.

**Fig S3.**
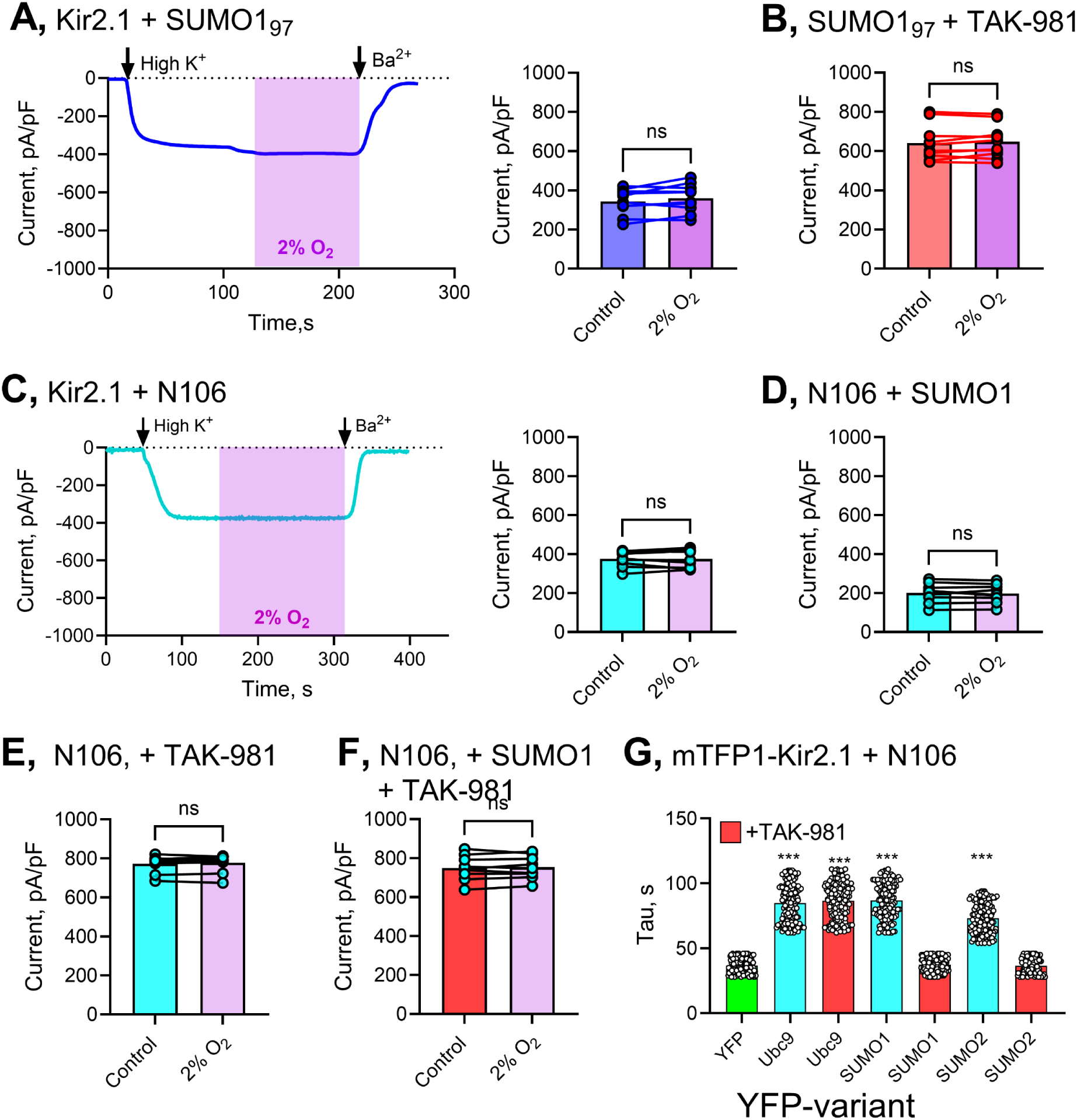
TAK-981 pretreatment prevents regulation of Kir2.1 by direct SUMOylation. FRET data are donor (mTFP) photobleaching curves, fit with a mono-exponential function to yield a fluorescence decay time constant (tau). Mean taus represent 80-120 regions of interest per group across at least 3 biological replicates. Patch-clamp data are from 6-12 cells per group. Kir2.1 currents were measured at -80 mV. Hypoxia is a drop in O_2_ from ambient levels to 2% measured at the cell. Purified SUMO1_97_ protein was applied at 1 µM in the recording pipette. TAK-981 was used at 100 nM and N106 was used at 10 µM. Each was applied 3 hours before the experiment. Statistical differences were determined using a paired, two-tailed Student’s t test where *** is p< 0.001. **A,** *Left*, Representative time-courses showing that Kir2.1 currents are suppressed and are insensitive to hypoxia when SUMO1_97_ is included in the recording pipette. *Right*, Summary histogram of paired recordings. **B,** Pretreatment with TAK-981 prevented the suppression of Kir2.1 current by purified SUMO1_97_. **C,** *Left*, Representative time-courses showing that Kir2.1 currents are suppressed and are insensitive to hypoxia following pretreatment with N106. *Right*, Summary histogram of paired recordings. **D,** Pretreatment with N106 enhances the suppression of Kir2.1 current evoked by purified SUMO1_97_. **E,** Suppression of Kir2.1 by N106 is prevented by TAK-981. **F,** Suppression of Kir2.1 by N106 and SUMO1_97_ is prevented by TAK-981. **G,** Summary histogram of photobleaching tau’s showing FRET between mTFP- tagged Kir2.1 and YFP-tagged Ubc9, SUMO1, or SUMO2 in the presence of N106. TAK-981 precludes FRET between the channel and SUMO1 or SUMO2 but not Ubc9,.

**Fig S4.**
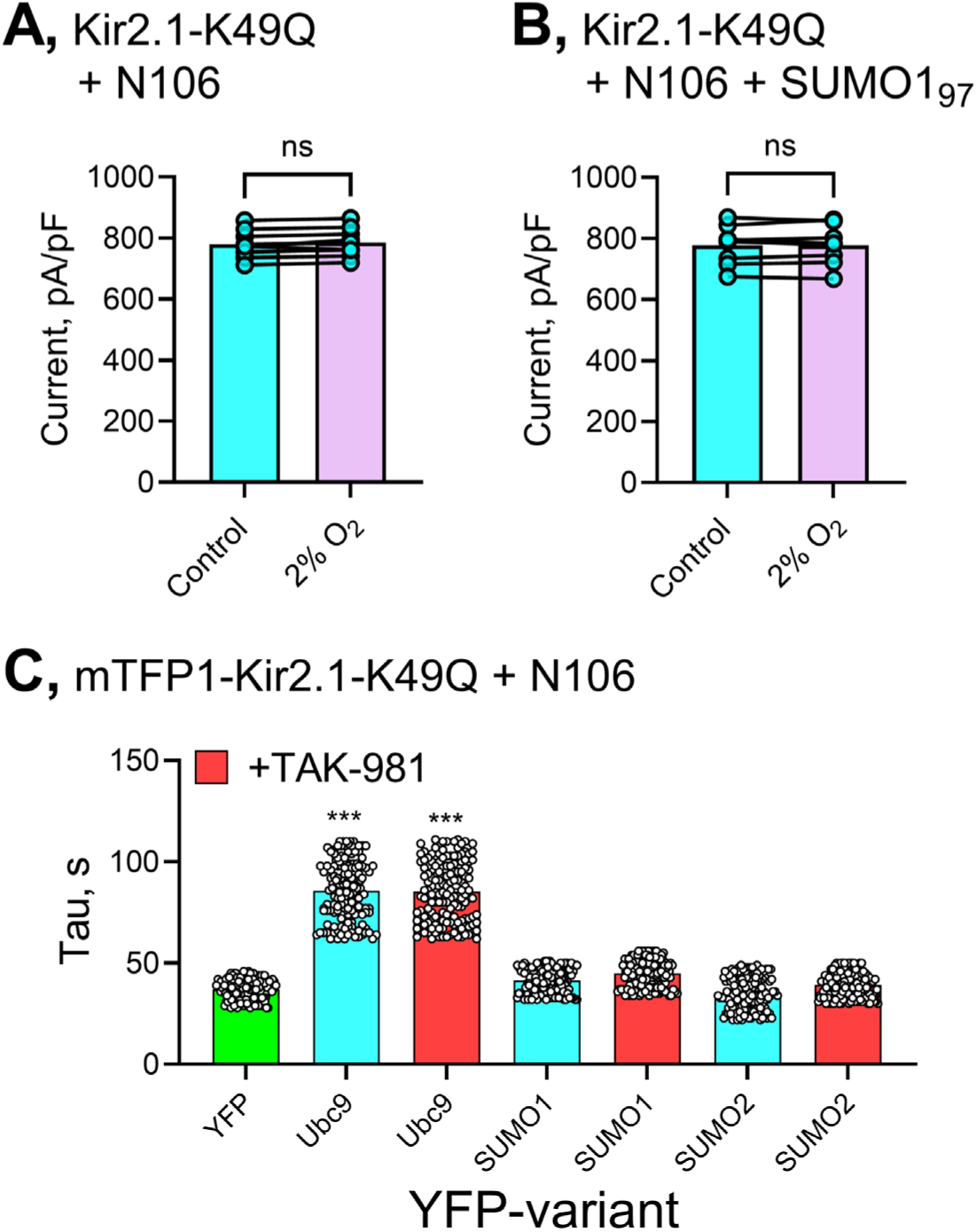
Kir2.1-K49Q channels are insensitive to N106. FRET data are donor (mTFP) photobleaching curves, fit with a mono-exponential function to yield a fluorescence decay time constant (tau). Mean taus represent 80-120 regions of interest per group across at least 3 biological replicates. Patch-clamp data are from 8-10 cells per group. Kir2.1 currents were measured at -80 mV. Hypoxia is a drop in O_2_ from ambient levels to 2% measured at the cell. Purified SUMO1_97_ protein was applied at 1 µM in the recording pipette. N106 was used at 10 µM and was applied 3 hours before the experiment. Statistical differences were determined using a paired, two-tailed Student’s t test where *** is p< 0.001. **A,** Summary histogram of paired recordings showing that N106 does not regulate Kir2.1-K49Q channels studied under ambient-O_2_ or hypoxic conditions. **B,** Summary histogram of paired recordings showing that N106 does not regulate Kir2.1-K49Q channels studied under ambient-O_2_ or hypoxic conditions with SUMO 1_97_ in the recording pipette. **C,** Summary histogram of photobleaching tau’s showing FRET between mTFP-tagged Kir2.1-K49Q and YFP-tagged Ubc9 but not SUMO1, or SUMO2 in the presence of N106.

**Fig S5.**
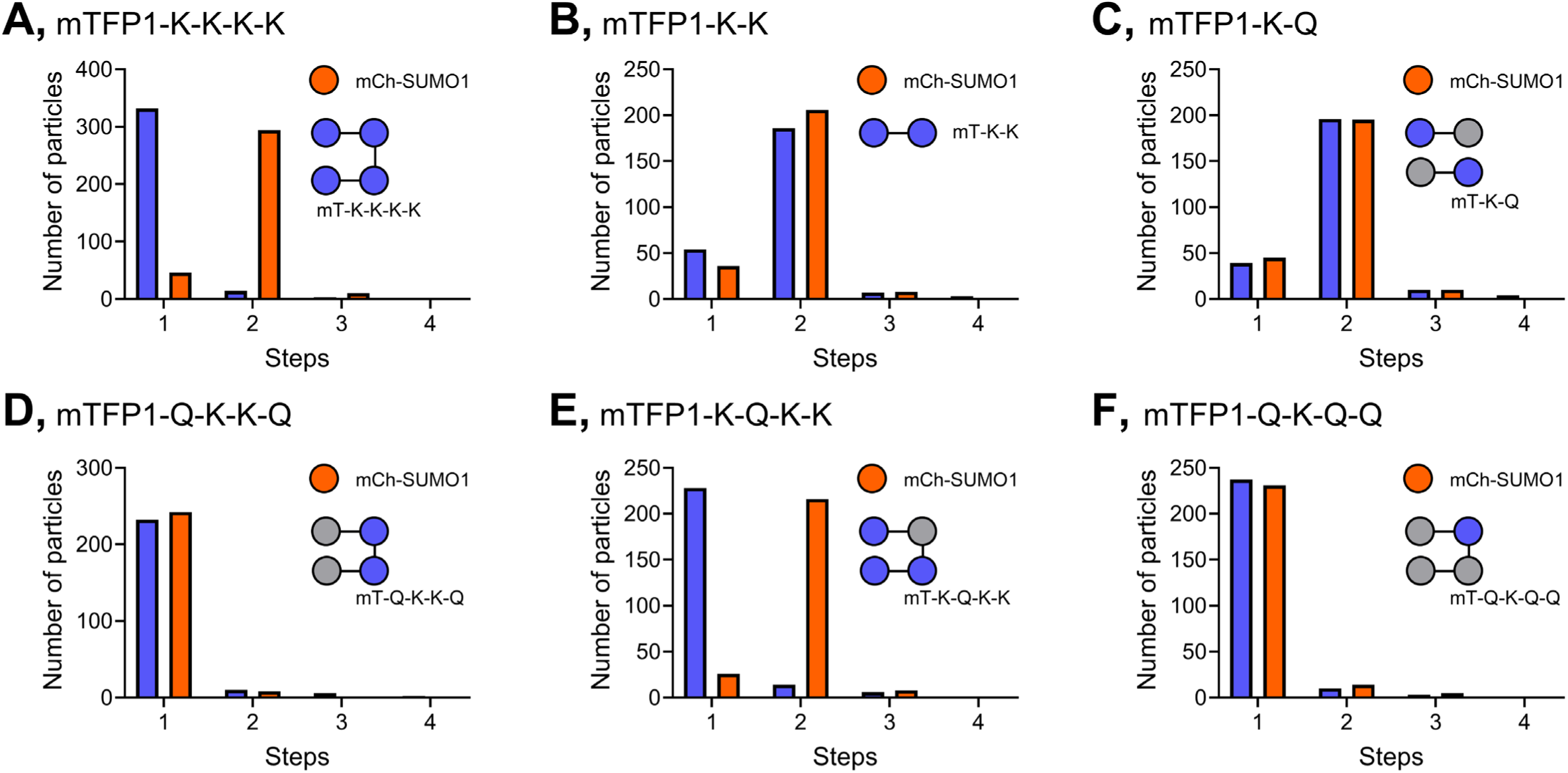
Kir2.1 channels are modified by up to two SUMO1 monomers. mCherry-SUMO1 (orange) was expressed with mTFP1-tagged Kir2.1 concatenated constructs in HEK293T cells and visualized at the membrane by TIRF microscopy. The stoichiometry of single Kir2.1-SUMO1 complexes was measured by single particle photobleaching, as per Figure 2. Wild type Kir2.1 subunits are shown in blue and denoted as K. Subunits with the K49Q mutation that cannot be SUMOylated are shown in gray and denoted as Q. **A,** Single fluorescent particles with a channel formed by one tetramer of wildtype Kir2.1 subunits (mTFP-K-K-K-K) associates with 2 mCherry-SUMO1s. **B,** Single fluorescent particles with a channel formed by two dimers of wildtype Kir2.1 subunits (mTFP-K-K) associates with 2 mCherry-SUMO1s. **C,** Single fluorescent particles with a channel formed by two dimers of wildtype and K49Q Kir2.1 subunits (mTFP-K-Q) associates with 2 mCherry-SUMO1s. **D,** Single fluorescent particles with a channel formed by one mTFP-Q-K-K-Q tetramer associates with 1 mCherry-SUMO1. **E,** Single fluorescent particles with a channel formed by one mTFP-K-Q-K-Q tetramer associates with 2 mCherry-SUMO1s. **F,** Single fluorescent particles with a channel formed by one mTFP-Q-K-Q-Q tetramer associates with 1 mCherry-SUMO1.

**Fig S6.**
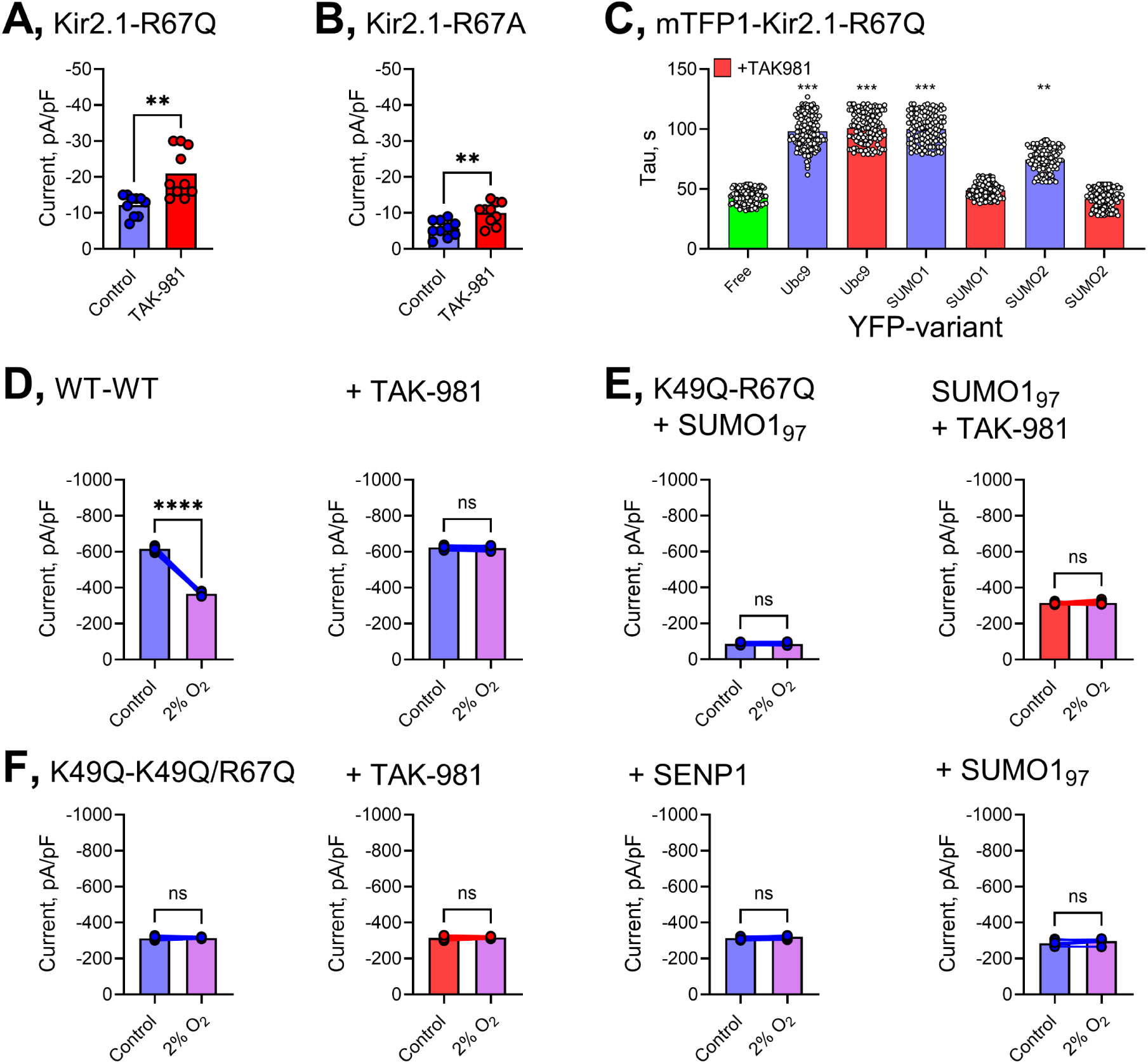
Characterization of Kir2.1-R67X monomers and dimers. FRET data donor (mTFP) are photobleaching curves, fit with a mono-exponential function to yield a fluorescence decay time constant (tau). Mean taus represent 80-120 regions of interest per group across at least 3 biological replicates. Patch-clamp data are from 9-14 cells per group. Kir2.1 currents were measured at -80 mV. Hypoxia is a drop in O_2_ from ambient levels to 2% measured at the cell. Purified SUMO1_97_ or SENP1 protein were applied at 1 µM and 2 µM, respectively, in the recording pipette. TAK-981 was used at 100 nM and was applied 3 hours before the experiment. Statistical differences were determined using two-tailed Student’s t tests where ** is P<0.01, *** is p< 0.001 and **** is P<0.0001 **A,** Summary histogram of Kir2.1-R67Q channels recorded with or without pretreatment with TAK-981. Summary histogram of Kir2.1-R67A channels recorded with or without pretreatment with TAK-981 **C,** Summary histogram of photobleaching tau’s showing FRET between mTFP-tagged Kir2.1- R67Q and YFP-tagged Ubc9, SUMO1 or SUMO2. FRET between the channel and SUMO1 or SUMO2 is abolished by pretreatment with TAK-981. **D,** Summary histograms of Kir2.1 channels formed by dimers (WT-WT) showing that hypoxic inhibition of the current is prevented by pretreatment with TAK-981 (right). **E,** Summary histogram of channels formed by dimers of Kir2.1-K49Q and Kir2.1-R67Q subunits showing that the current is suppressed when 1 µM SUMO1_97_ is included in the recording pipette. **F,** Summary histograms of channels formed by dimers of subunits with Kir2.1-K49Q and the double mutant subunit Kir2.1-K49Q/R67Q subunits. The currents are not modulated by hypoxia, by pretreatment with TAK-981, or when 1 µM SUMO1_97_, or 2 µM SENP1 were included in the recording pipette.

## References

1. Handklo-Jamal R, Meisel E, Yakubovich D, Vysochek L, Beinart R, Glikson M, McMullen JR, Dascal N, Nof E and Oz S. Andersen-Tawil Syndrome Is Associated With Impaired PIP(2) Regulation of the Potassium Channel Kir2.1. Front Pharmacol. 2020;11:672.

2. Tawil R, Ptacek LJ, Pavlakis SG, DeVivo DC, Penn AS, Ozdemir C and Griggs RC. Andersen’s syndrome: potassium-sensitive periodic paralysis, ventricular ectopy, and dysmorphic features. Ann Neurol. 1994;35:326–30.

3. Ptacek LJ, George AL, Jr., Griggs RC, Tawil R, Kallen RG, Barchi RL, Robertson M and Leppert MF. Identification of a mutation in the gene causing hyperkalemic periodic paralysis. Cell. 1991;67:1021–7.

4. Donaldson MR, Jensen JL, Tristani-Firouzi M, Tawil R, Bendahhou S, Suarez WA, Cobo AM, Poza JJ, Behr E, Wagstaff J, Szepetowski P, Pereira S, Mozaffar T, Escolar DM, Fu YH and Ptacek LJ. PIP2 binding residues of Kir2.1 are common targets of mutations causing Andersen syndrome. Neurology. 2003;60:1811–6.

5. Plaster NM, Tawil R, Tristani-Firouzi M, Canun S, Bendahhou S, Tsunoda A, Donaldson MR, Iannaccone ST, Brunt E, Barohn R, Clark J, Deymeer F, George AL, Jr., Fish FA, Hahn A, Nitu A, Ozdemir C, Serdaroglu P, Subramony SH, Wolfe G, Fu YH and Ptacek LJ. Mutations in Kir2.1 cause the developmental and episodic electrical phenotypes of Andersen’s syndrome. Cell. 2001;105:511–9.

6. Perez-Riera AR, Barbosa-Barros R, Samesina N, Pastore CA, Scanavacca M, Daminello- Raimundo R, de Abreu LC, Nikus K and Brugada P. Andersen-Tawil Syndrome: A Comprehensive Review. Cardiol Rev. 2021;29:165–177.

7. Zhang L, Benson DW, Tristani-Firouzi M, Ptacek LJ, Tawil R, Schwartz PJ, George AL, Horie M, Andelfinger G, Snow GL, Fu YH, Ackerman MJ and Vincent GM. Electrocardiographic features in Andersen-Tawil syndrome patients with KCNJ2 mutations: characteristic T-U-wave patterns predict the KCNJ2 genotype. Circulation. 2005;111:2720–6.

8. Sansone V and Tawil R. Management and treatment of Andersen-Tawil syndrome (ATS). Neurotherapeutics. 2007;4:233–7.

9. Jeevaratnam K, Chadda KR, Salvage SC, Valli H, Ahmad S, Grace AA and Huang CL. Ion channels, long QT syndrome and arrhythmogenesis in ageing. Clin Exp Pharmacol Physiol. 2017;44 Suppl 1:38–45.

10. Ayres SM and Grace WJ. Inappropriate ventilation and hypoxemia as causes of cardiac arrhythmias. The control of arrhythmias without antiarrhythmic drugs. Am J Med. 1969;46:495–505.

11. Plant LD, Xiong D, Romero J, Dai H and Goldstein SAN. Hypoxia Produces Pro-arrhythmic Late Sodium Current in Cardiac Myocytes by SUMOylation of NaV1.5 Channels. Cell Rep. 2020;30:2225–2236 e4.

12. Keating MT and Sanguinetti MC. Molecular and cellular mechanisms of cardiac arrhythmias. Cell. 2001;104:569–80.

13. Bezzina CR, Lahrouchi N and Priori SG. Genetics of sudden cardiac death. Circ Res. 2015;116:1919–36.

14. Dhamoon AS and Jalife J. The inward rectifier current (IK1) controls cardiac excitability and is involved in arrhythmogenesis. Heart Rhythm. 2005;2:316–24.

15. Xu Y, Yang Y, Chandrashekar A, Gada KD, Masotti M, Baggetta AM, Connolly JG, Kawano T and Plant LD. Hypoxia inhibits the cardiac I K1 current through SUMO targeting Kir2.1 activation by PIP2. iScience. 2022;25:104969.

16. Ackerman MJ and Giudicessi JR. Time to Redefine the Natural History and Clinical Management of Type 1 Andersen-Tawil Syndrome? J Am Coll Cardiol. 2020;75:1785–1787.

17. Logothetis DE, Petrou VI, Zhang M, Mahajan R, Meng XY, Adney SK, Cui M and Baki L. Phosphoinositide control of membrane protein function: a frontier led by studies on ion channels. Annu Rev Physiol. 2015;77:81–104.

18. Lopes CM, Zhang H, Rohacs T, Jin T, Yang J and Logothetis DE. Alterations in conserved Kir channel-PIP2 interactions underlie channelopathies. Neuron. 2002;34:933–44.

19. Liang S, Wang Q, Zhang W, Zhang H, Tan S, Ahmed A and Gu Y. Carbon monoxide inhibits inward rectifier potassium channels in cardiomyocytes. Nat Commun. 2014;5:4676.

20. Connolly JG and Plant LD. SUMO Regulation of Ion Channels in Health and Disease. Physiology (Bethesda). 2025;40:0.

21. Henley JM, Carmichael RE and Wilkinson KA. Extranuclear SUMOylation in Neurons. Trends Neurosci. 2018;41:198–210.

22. Jansen LR, Welch MA, Plant LD and Baro DJ. Crosstalk between PKA and PIAS3 regulates cardiac Kv4 channel SUMOylation. Cell Commun Signal. 2024;22:422.

23. Edelstein AD, Tsuchida MA, Amodaj N, Pinkard H, Vale RD and Stuurman N. Advanced methods of microscope control using muManager software. J Biol Methods. 2014;1.

24. Schneider CA, Rasband WS and Eliceiri KW. NIH Image to ImageJ: 25 years of image analysis. Nat Methods. 2012;9:671–5.

25. Gordon JC, Myers JB, Folta T, Shoja V, Heath LS and Onufriev A. H++: a server for estimating pKas and adding missing hydrogens to macromolecules. Nucleic Acids Res. 2005;33:W368–71.

26. Leal-Pinto E, London RD, Knorr BA and Abramson RG. Reconstitution of hepatic uricase in planar lipid bilayer reveals a functional organic anion channel. J Membr Biol. 1995;146:123–32.

27. Jo S, Kim T, Iyer VG and Im W. CHARMM-GUI: a web-based graphical user interface for CHARMM. J Comput Chem. 2008;29:1859–65.

28. Case DA, Cheatham TE, 3rd, Darden T, Gohlke H, Luo R, Merz KM, Jr., Onufriev A, Simmerling C, Wang B and Woods RJ. The Amber biomolecular simulation programs. J Comput Chem. 2005;26:1668–88.

29. Li D, Jin T, Gazgalis D, Cui M and Logothetis DE. On the mechanism of GIRK2 channel gating by phosphatidylinositol bisphosphate, sodium, and the Gbetagamma dimer. J Biol Chem. 2019;294:18934–18948.

30. Breneman CM and Wiberg KB. Determining Atom-Centered Monopoles from Molecular Electrostatic Potentials - the Need for High Sampling Density in Formamide Conformational- Analysis. Journal of Computational Chemistry. 1990;11:361–373.

31. Darden T, York D and Pedersen L. Particle mesh Ewald: An N⋅log(N) method for Ewald sums in large systems. The Journal of Chemical Physics. 1993;98:10089–10092.

32. Langston SP, Grossman S, England D, Afroze R, Bence N, Bowman D, Bump N, Chau R, Chuang BC, Claiborne C, Cohen L, Connolly K, Duffey M, Durvasula N, Freeze S, Gallery M, Galvin K, Gaulin J, Gershman R, Greenspan P, Grieves J, Guo J, Gulavita N, Hailu S, He X, Hoar K, Hu Y, Hu Z, Ito M, Kim MS, Lane SW, Lok D, Lublinsky A, Mallender W, McIntyre C, Minissale J, Mizutani H, Mizutani M, Molchinova N, Ono K, Patil A, Qian M, Riceberg J, Shindi V, Sintchak MD, Song K, Soucy T, Wang Y, Xu H, Yang X, Zawadzka A, Zhang J and Pulukuri SM. Discovery of TAK-981, a First-in-Class Inhibitor of SUMO-Activating Enzyme for the Treatment of Cancer. J Med Chem. 2021;64:2501–2520.

33. Cotelle N, Bernier JL, Henichart JP, Catteau JP, Gaydou E and Wallet JC. Scavenger and antioxidant properties of ten synthetic flavones. Free Radic Biol Med. 1992;13:211–9.

34. Kim YS, Nagy K, Keyser S and Schneekloth JS, Jr. An electrophoretic mobility shift assay identifies a mechanistically unique inhibitor of protein sumoylation. Chem Biol. 2013;20:604–13.

35. Kotler O, Khrapunsky Y, Shvartsman A, Dai H, Plant LD, Goldstein SAN and Fleidervish I. SUMOylation of Na(V)1.2 channels regulates the velocity of backpropagating action potentials in cortical pyramidal neurons. Elife. 2023;12.

36. Xiong D, Li T, Dai H, Arena AF, Plant LD and Goldstein SAN. SUMOylation determines the voltage required to activate cardiac IKs channels. Proc Natl Acad Sci U S A. 2017;114:E6686–E6694.

37. Welch MA, Jansen LR and Baro DJ. SUMOylation of the Kv4.2 Ternary Complex Increases Surface Expression and Current Amplitude by Reducing Internalization in HEK 293 Cells. Front Mol Neurosci. 2021;14:757278.

38. Kho C, Lee A, Jeong D, Oh JG, Gorski PA, Fish K, Sanchez R, DeVita RJ, Christensen G, Dahl R and Hajjar RJ. Small-molecule activation of SERCA2a SUMOylation for the treatment of heart failure. Nat Commun. 2015;6:7229.

39. Plant LD, Marks JD and Goldstein SA. SUMOylation of NaV1.2 channels mediates the early response to acute hypoxia in central neurons. Elife. 2016;5.

40. Kalscheur MM, Vaidyanathan R, Orland KM, Abozeid S, Fabry N, Maginot KR, January CT, Makielski JC and Eckhardt LL. KCNJ2 mutation causes an adrenergic-dependent rectification abnormality with calcium sensitivity and ventricular arrhythmia. Heart Rhythm. 2014;11:885–94.

41. D’Avanzo N, Lee SJ, Cheng WWL and Nichols CG. Energetics and location of phosphoinositide binding in human Kir2.1 channels. J Biol Chem. 2013;288:16726–16737.

42. Fernandes CAH, Zuniga D, Fagnen C, Kugler V, Scala R, Pehau-Arnaudet G, Wagner R, Perahia D, Bendahhou S and Venien-Bryan C. Cryo-electron microscopy unveils unique structural features of the human Kir2.1 channel. Sci Adv. 2022;8:eabq8489.

43. Gada KD, Xu Y, Winn BT, Masotti M, Kawano T, Vaananen H and Plant LD. Optogenetic dephosphorylation of phosphatidylinositol 4,5 bisphosphate in Xenopus laevis oocytes. STAR Protoc. 2023;4:102003.

44. Ningoo M, Plant LD, Greka A and Logothetis DE. PIP2 regulation of TRPC5 channel activation and desensitization. J Biol Chem. 2021;296:100726.

45. Rohacs T, Chen J, Prestwich GD and Logothetis DE. Distinct specificities of inwardly rectifying K(+) channels for phosphoinositides. J Biol Chem. 1999;274:36065–72.

46. Ha J, Xu Y, Kawano T, Hendon T, Baki L, Garai S, Papapetropoulos A, Thakur G, Plant LD and Logothetis DE. Hydrogen sulfide inhibits Kir2 and Kir3 channels by decreasing sensitivity to the phospholipid PIP2. J Biol Chem. 2018.

47. Plant LD, Dowdell EJ, Dementieva IS, Marks JD and Goldstein SA. SUMO modification of cell surface Kv2.1 potassium channels regulates the activity of rat hippocampal neurons. J Gen Physiol. 2011;137:441–54.

48. Kharche S, Garratt CJ, Boyett MR, Inada S, Holden AV, Hancox JC and Zhang H. Atrial proarrhythmia due to increased inward rectifier current (I(K1)) arising from KCNJ2 mutation--a simulation study. Prog Biophys Mol Biol. 2008;98:186–97.

49. Sattler SM, Skibsbye L, Linz D, Lubberding AF, Tfelt-Hansen J and Jespersen T. Ventricular Arrhythmias in First Acute Myocardial Infarction: Epidemiology, Mechanisms, and Interventions in Large Animal Models. Front Cardiovasc Med. 2019;6:158.

50. Liebelt F, Sebastian RM, Moore CL, Mulder MPC, Ovaa H, Shoulders MD and Vertegaal ACO. SUMOylation and the HSF1-Regulated Chaperone Network Converge to Promote Proteostasis in Response to Heat Shock. Cell Rep. 2019;26:236–249 e4.

51. Zhou W, Ryan JJ and Zhou H. Global analyses of sumoylated proteins in Saccharomyces cerevisiae. Induction of protein sumoylation by cellular stresses. J Biol Chem. 2004;279:32262–8.

52. Wang Y, Gao Y, Tian Q, Deng Q, Wang Y, Zhou T, Liu Q, Mei K, Wang Y, Liu H, Ma R, Ding Y, Rong W, Cheng J, Yao J, Xu TL, Zhu MX and Li Y. TRPV1 SUMOylation regulates nociceptive signaling in models of inflammatory pain. Nat Commun. 2018;9:1529.

53. Lightcap ES, Yu P, Grossman S, Song K, Khattar M, Xega K, He X, Gavin JM, Imaichi H, Garnsey JJ, Koenig E, Zhang H, Lu Z, Shah P, Fu Y, Milhollen MA, Hatton BA, Riceberg J, Shinde V, Li C, Minissale J, Yang X, England D, Klinghoffer RA, Langston S, Galvin K, Shapiro G, Pulukuri SM, Fuchs SY and Huszar D. A small-molecule SUMOylation inhibitor activates antitumor immune responses and potentiates immune therapies in preclinical models. Sci Transl Med. 2021;13:eaba7791.

54. Derry JMJ, Burns C, Frazier JP, Beirne E, Grenley M, DuFort CC, Killingbeck E, Leon M, Williams C, Gregory M, Houlton J, Clayburgh D, Swiecicki P, Huszar D, Berger A and Klinghoffer RA. Trackable Intratumor Microdosing and Spatial Profiling Provide Early Insights into Activity of Investigational Agents in the Intact Tumor Microenvironment. Clin Cancer Res. 2023;29:3813–3825.

